# Experimental evolution of *Bacillus subtilis* on *Arabidopsis thaliana* roots reveals fast adaptation and improved root colonization in the presence of soil microbes

**DOI:** 10.1101/2021.07.09.451762

**Authors:** Mathilde Nordgaard, Christopher Blake, Gergely Maróti, Guohai Hu, Yue Wang, Mikael Lenz Strube, Ákos T. Kovács

## Abstract

The soil ubiquitous *Bacillus subtilis* is known to promote plant growth and protect plants against disease. These characteristics make *B. subtilis* highly relevant in an agricultural perspective, fueling the interest in studying *B. subtilis*-plant interactions. Here, we employ an experimental evolution approach to explore adaptation of *B. subtilis* to *Arabidopsis thaliana* roots. We found that *B. subtilis* rapidly adapted to the plant root environment, as evidenced by improved root colonizers observed already after 12 consecutive transfers between seedlings in a hydroponic setup. In addition, two selected evolved isolates from independent populations from transfer 30 outcompeted the ancestor during root colonization. Re-sequencing of single evolved isolates and endpoint populations revealed mutations in genes related to different bacterial traits. Further, phenotypic characterization of evolved isolates from transfer 30 showed that increased root colonization was associated with robust biofilm formation in response to the plant polysaccharide xylan. Additionally, several evolved isolates across independent populations were impaired in motility, a redundant trait in the selective environment. Interestingly, two evolved isolates suffered a fitness disadvantage in a non-selective environment, demonstrating an evolutionary cost of adaptation to the plant root. Finally, increased root colonization by a selected evolved isolate was also demonstrated in the presence of resident soil microbes. Our findings provide novel insights into how a well-known plant growth-promoting rhizobacterium rapidly adapts to an ecologically relevant environment and reveal evolutionary consequences that are fundamental to consider when evolving strains for biocontrol purposes.

## Introduction

The nutrient-rich rhizosphere is a hotspot for microbial activity, containing up to 10^11^ bacteria per gram root (Egamberdieva et al., 2008) and housing more than 30.000 prokaryotic species (Mendes et al., 2011). Among those are beneficial bacteria which are actively recruited by the plant through root exudate secretion and subsequently colonize the root from where they benefit the plant through various mechanisms (Berendsen et al., 2018, 2012; Mendes et al., 2011; Rudrappa et al., 2008; Trivedi et al., 2020). One well-known plant growth-promoting rhizobacterium (PGPR) is the spore-forming *Bacillus subtilis* which has been isolated from various plant species (Cazorla et al., 2007; Fall et al., 2004; Huang et al., 2017; Pandey and Palni, 1997). Its plant-beneficial traits (Blake et al., 2021a) and promising role as a biocontrol agent (Fira et al., 2018; Kiesewalter et al., 2021; Ongena and Jacques, 2008) have fueled the interest in studying *B. subtilis*-plant interactions and led to the elucidation of mechanisms involved in the establishment of *B. subtilis* on the root and behind its plant beneficial properties.

An obvious prerequisite for successful root colonization is the ability of the bacterium to reach the plant root. Chemotaxis towards root exudates was shown to be important for the early colonization of *Arabidopsis thaliana* roots by *B. subtilis* under hydroponic conditions (Allard-Massicotte et al., 2016), while solid surface motility has been suggested to play a role during tomato root colonization in vermiculites (Tian et al., 2021). After reaching the plant root, *B. subtilis* initiates biofilm formation (Allard-Massicotte et al., 2016; Bais et al., 2004; Beauregard et al., 2013; Chen et al., 2013). Similar to *in vitro* conditions, the formation of plant root-associated biofilms depends on the production of the matrix components EPS and TasA (Beauregard et al., 2013; Branda et al., 2006; Chen et al., 2013; Dragoš et al., 2018a). Expression of the operons involved in matrix production, *epsA-O* and *tapA-sipW-tasA,* is controlled by the biofilm repressor SinR (Chu et al., 2006; Kearns et al., 2005). In response to environmental cues, one or more of the five histidine kinases, KinA-E, are activated resulting in phosphorylation of the master regulator Spo0A through a phosphorelay (Jiang et al., 2000). At threshold concentrations of Spo0A∼P, SinI is produced (Fujita et al., 2005), which binds to and inhibits SinR (Bai et al., 1993), resulting in matrix gene expression. As an environmental cue for root colonization, root exudates from tomato were shown to trigger biofilm formation in *B. subtilis* in a KinD-dependent manner (Chen et al., 2012), while another study reported biofilm induction by plant polysaccharides via KinC and KinD (Beauregard et al., 2013). *B. subtilis* benefits from such microbe-plant interactions by acquiring carbon source from the plants. In turn, *B. subtilis* protects the plant against disease directly by producing antimicrobials (Asaka and Shoda, 1996; Bais et al., 2004; Chen et al., 2013; Kiesewalter et al., 2021) and indirectly through niche competition (Köhl et al., 2019; Lugtenberg and Kamilova, 2009) and elicitation of induced systemic resistance in the plant (Akram et al., 2015; Rudrappa et al., 2008). Moreover, *B. subtilis* promotes plant growth by improving nutrient availability and producing growth-promoting phytohormones (Blake et al., 2021a).

Mutualistic bacteria-plant interactions are a result of a long-term co-evolution of bacteria and plants that started by the colonization of land by ancestral plants 450 million years ago (Hassani et al., 2018). Here, we were interested in studying how *B. subtilis* adapts to plant roots on a much shorter evolutionary time scale. Experimental evolution (EE) provides a powerful tool to study microbial adaptation to different environments in real-time (Kawecki et al., 2012; Lenski, 2017). We recently studied EE of *B. subtilis* on *A. thaliana* plants roots, which revealed diversification of *B. subtilis* into three distinct morphotypes. A mix of the three morphotypes displayed increased root colonization compared to the sum of the three morphotypes in monocultures weighted by their initial relative abundance in the mix, which was demonstrated to be caused by complementarity effects (Blake et al., 2021b). Such morphological diversification has also been observed in EE of *B. subtilis* biofilm pellicles formed at the air-liquid interface (Dragoš et al., 2018b) as well as during EE of *Burkholderia cenocepacia* biofilms on polystyrene beads (Poltak and Cooper, 2011). In this study, we performed EE of *B. subtilis* on one-week old *A. thaliana* roots under axenic conditions with the initial hypothesis that *B. subtilis* would adapt to the plant root environment by acquiring mutations that would provide the bacteria with a fitness advantage over the ancestor during root colonization. We found that *B. subtilis* rapidly adapted to the plant root as observed by improved root colonizers already after 12 consecutive transfers. In addition, two selected evolved isolates from independent populations from the final transfer (transfer 30) outcompeted the ancestor during root colonization. Furthermore, re-sequencing of single evolved isolates from independent populations and different time points revealed that the evolved isolates had acquired mutations in genes related to different bacterial traits. To further elucidate which bacterial traits were altered during the adaptation to plant roots, evolved isolates from the final transfer were subjected to additional phenotypic characterization. We found that evolved isolates from independent populations displayed robust biofilm formation in response to plant polysaccharides, impaired motility, and altered growth on plant compounds. Finally, we demonstrate that adaptation of *B. subtilis* to *A. thaliana* roots is accompanied by an evolutionary cost, and report an evolved isolate displaying increased root colonization also in the presence of resident soil microbes.

## Results

### *B. subtilis* populations evolved on *A. thaliana* roots show rapid increase in root colonization

To explore the evolutionary adaptation of *B. subtilis* to plant roots, we employed an experimental evolution (EE) setup previously established for another *Bacillus* species (Lin et al., 2021). In short, *B. subtilis* DK1042 (hereafter referred to as the ancestor) was inoculated onto *A. thaliana* seedlings under hydroponic conditions in seven parallel populations. The MSNg medium used in the EE is a minimal medium supplemented with a very low concentration of glycerol (0.05 %), and the bacteria thereby move towards the plant root to access a carbon source. Every 48 h for a total of 64 days, the newly colonized seedling was transferred to fresh medium containing a new sterile seedling, thereby enabling re-colonization (Fig. 1A). We hypothesized, that during the successive transfers *B. subtilis* would adapt to the plant roots by acquiring mutations that would confer a fitness advantage over the ancestor during root colonization, resulting in these mutations being selected for. In this setup, we specifically selected for a regular cycle of dispersal from the root-associated biofilm, chemotaxis towards the new root and biofilm formation on the root surface. To follow potential changes in root colonization of the evolving populations during the ongoing EE, the productivity, i.e. colony-forming unit (CFU) per root, was quantified at different time points. All seven independent populations showed a rapid increase in root colonization within the first seven transfers, after which the productivity of the populations remained rather stable with slight increases and drops dependent on the certain population (Fig. 1B). While this rapid increase in productivity could be due to genetic adaptation to the plant root, the initial rise could also be caused by physiological adaptation to the experimental conditions. Interestingly, such a rapid increase in productivity of populations evolving on plant roots is consistent with our recent study where *B. subtilis* was evolved on older *A. thaliana* roots (Blake et al., 2021b).

**Fig. 1:**
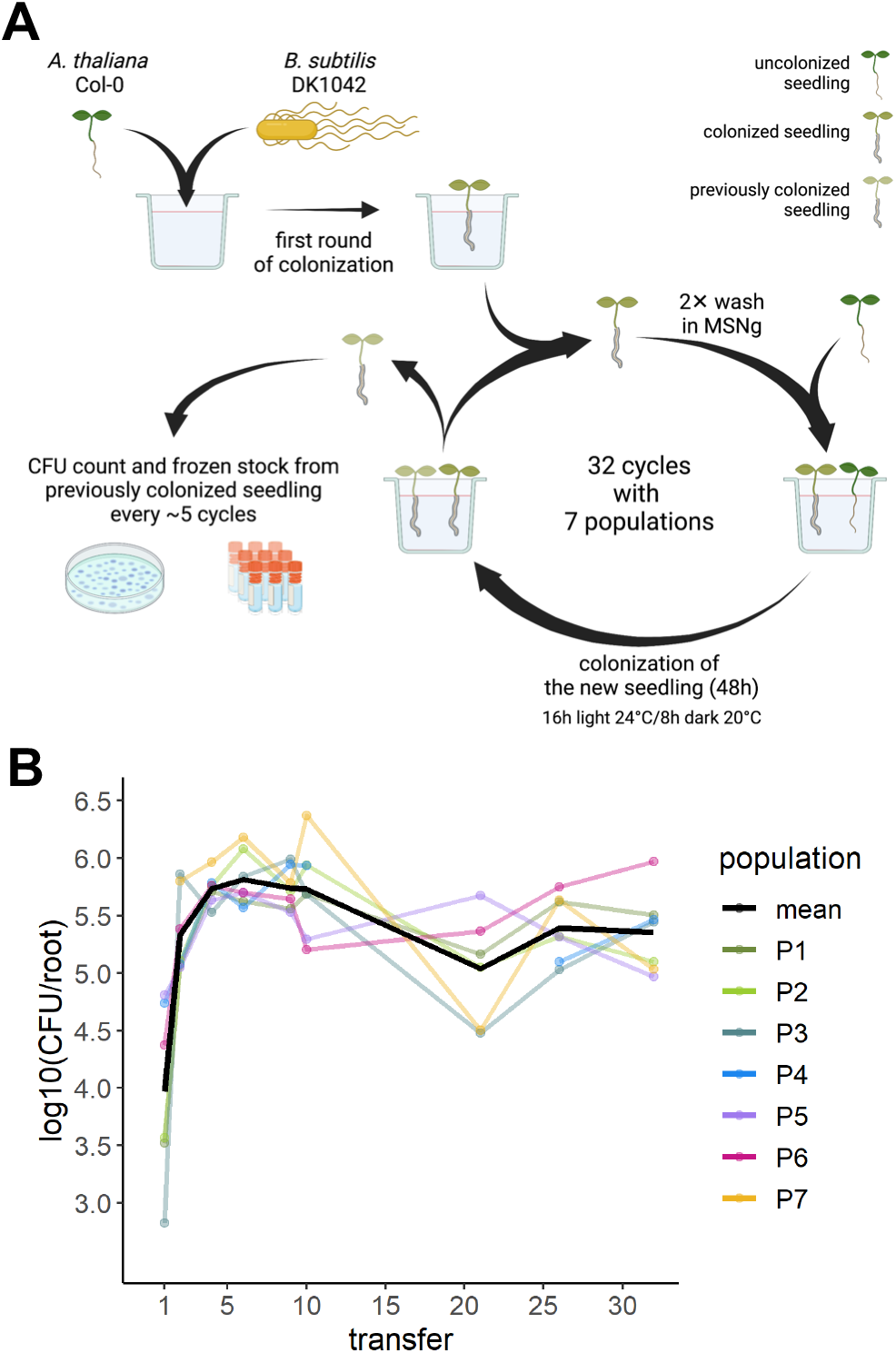
Overview of experimental evolution and productivity of evolving populations. (**A**) Overview of the experimental evolution approach. (**B**) The *B. subtilis* populations rapidly increased in productivity during the experimental evolution on *A. thaliana* roots. The productivity (CFU/root) of the evolving populations was systematically quantified as CFU/root at 9 different time points during the ongoing EE. The y-axis displays the log10-transformed productivity. The black line represents the mean population productivity (N=7).

### Several evolved isolates display altered colony morphologies

To examine whether genetic adaptation to the plant root had taken place during the EE, single evolved isolates from the evolved populations were saved as frozen stocks and subjected to phenotypic and genotypic characterization. To represent different populations and time points during the EE, three isolates were randomly picked from each of population 3, 4, 6 and 7 at transfer 12, 18 and 30 (hereafter referred to as T12, T18 and T30). To detect possible changes in colony morphology, ON cultures of the ancestor and evolved isolates were spotted on LB agar and colonies inspected after 48 h incubation. On LB agar, the ancestor formed a round colony with a wrinkled periphery, while different colony morphologies were observed among the evolved isolates (Fig. 2). At T12, some isolates displayed a colony morphology resembling the ancestor, e.g. isolate 3 from population 3 (3.3) and isolate 2 from population 7 (7.2), referred to as the “Wrinkled”-type. Several other isolates formed a colony with a white sharp edge along the wrinkled periphery (including isolates 3.2, 4.3, 6.1 and 7.3), hereafter referred to as the “Sharp-Wrinkled”-type. Additionally, isolate 7.1 formed a hyper-wrinkled, white colony, referred to as the “Snow”-type. These distinct colony morphologies were also observed at later time points (T18 and T30). Interestingly, the Snow-type was only observed in population 7. Furthermore, the three isolates from population 6 at T30 formed slightly less wrinkled colonies compared to the ancestor. We note that three isolates do not represent the entire population, and isolates with other colony morphologies could be present in the populations. Nonetheless, the appearance of isolates with altered colony morphologies in the four populations indicates the presence of genetic changes. Furthermore, the occurrence of isolates with altered colony morphologies already at T12, and especially the presence of three different types (Wrinkled, Sharp-Wrinkled and Snow) in population 7, at this early time point, suggests rapid diversification of *B. subtilis* during EE on *A. thaliana* roots. Such diversification into distinct morphotypes was also observed in our previous study on EE of *B. subtilis* on plant roots (Blake et al., 2021b) and has additionally been observed in EE of *B. subtilis* pellicle biofilms (Dragoš et al., 2018b), indicating successful adaptation to the selective environment.

**Fig. 2:**
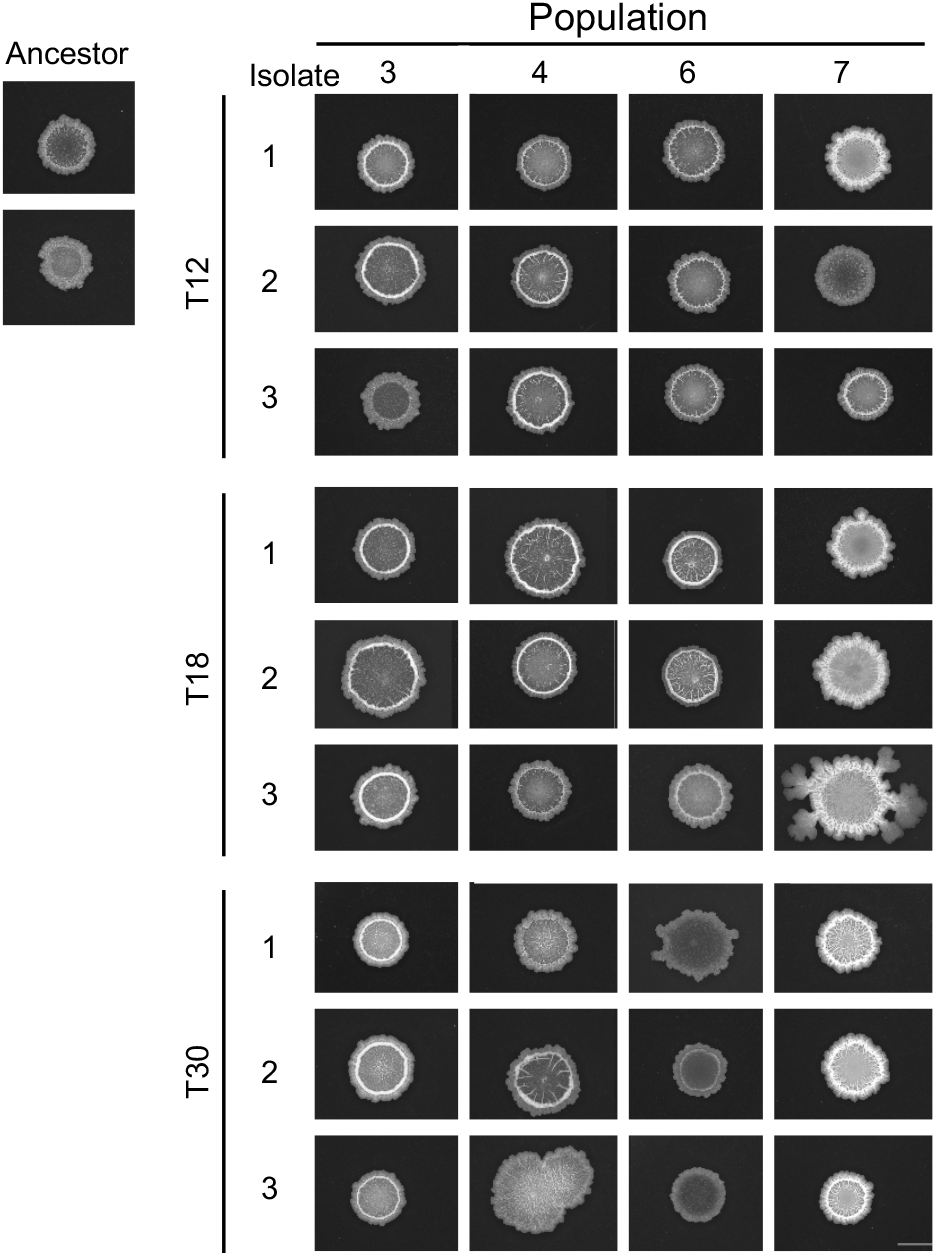
Distinct colony morphologies are observed among evolved isolates from different time points of the experimental evolution. ON cultures of the ancestor and evolved isolates from populations 3, 4, 6 and 7 at transfer 12, 18 and 30 were spotted on LB agar (1.5 %) and imaged after incubation for 48 h at 30 °C using a stereomicroscope. Ancestor represents *B. subtilis* DK1042. Each colony is representative of at least three replicates. Scale bar = 5 mm.

### Evolved isolates from different time points show increased colonization of *A. thaliana* roots

The design of the EE employed in this study should enable selection for bacteria that efficiently colonize the root. We, therefore, speculated whether the altered colony morphology of some of the evolved isolates was associated with improved productivity on the root (CFU/mm root). To test this, the ancestor and evolved isolates from the final time point (T30) were tested for individual colonization of *A. thaliana* seedlings under the same conditions applied during the EE. CFU quantification revealed that most evolved isolates tended to show increased root colonization, with five isolates from three different populations displaying significantly increased productivity on the root, with an up to circa 1.3-fold change relative to the ancestor (Fig. 3). To track down when such improved root colonizers emerged during the EE, the randomly selected evolved isolates from T12 and T18 were similarly tested. Three and five evolved isolates at T12 and T18, respectively, displayed significantly increased productivity relative to the ancestor. In addition, a single isolate from T12 was significantly reduced in root colonization. These results confirm that indeed genetic adaptation of *B. subtilis* to the plant root took place during the EE. Furthermore, the observation of improved root colonizers already at T12 indicates that *B. subtilis* rapidly adapted to plant root colonization during the EE.

**Fig. 3:**
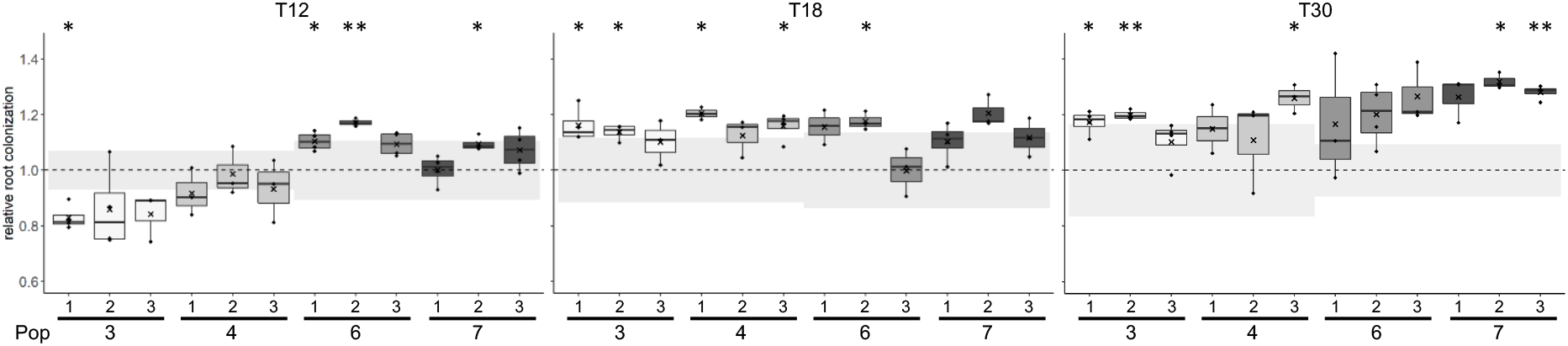
Evolved isolates from different time points show increased productivity on the root relative to the ancestor. The ancestor and evolved isolates from populations 3, 4, 6 and 7 at three different time points (T12, 18, 30) were tested for individual root colonization. For each evolved isolate, relative root colonization was calculated by dividing the log10-transformed productivity (CFU/mm root) of each replicate by the mean of the log10-transformed productivity of the ancestor from the same experimental setup. The cross represents the mean relative root colonization (N=3-4). The dashed, horizontal line represents the mean of the ancestor (N=3-4), while the grey-shaded rectangles represent the standard deviation of the ancestor from the corresponding experiment. The normalized values were subjected to a One-sample *t*-test to test whether the mean was significantly different from 1. P-values have been corrected for multiple comparisons. *: P<0.05, **: P<0.01.

### Selected evolved isolates display a fitness advantage over the ancestor that is specific to the plant root environment

While multiple evolved isolates displayed increased individual root colonization relative to the ancestor (Fig. 3), we next wanted to test whether the evolved isolates had a noticeable fitness advantage over the ancestor during competition on the root. For this purpose, two selected evolved isolates from independent populations from T30 (Ev6.1 and Ev7.3, referring to isolate 1 from population 6 and isolate 3 from population 7, respectively) were competed against the ancestor on the plant root. Following 48 h of root colonization, CFU quantification revealed that both evolved isolates had outcompeted the ancestor on the root, and statistical analysis confirmed that the evolved isolates had a significantly higher fitness relative to the ancestor (Fig. 4A; for calculation of relative fitness, see STAR Methods). This result was further supported by CLSM imaging: regardless of the fluorescence labeling combination, the two evolved isolates formed biofilms on the roots, as evidenced by aggregates along the root, while the ancestor was scarcely present (Fig. 4C). Noticeably, the fluorescent images revealed that Ev6.1 formed fewer and smaller aggregates along the root compared to Ev7.3 (Fig. 4C), consistent with the individual root colonization of the two isolates (Fig. 3). To test whether the fitness advantage of Ev6.1 and Ev7.3 over the ancestor was specific to the plant root environment, the two evolved isolates were competed against the ancestor in a non-selective environment, i.e. in LB supplemented with xylan, a plant polysaccharide (PP) found among others in the secondary cell walls of *A. thaliana* (Liepman et al., 2010), and under well-shaking conditions. In this non-selective environment, neither of the evolved isolates outcompeted the ancestor but instead seemed to suffer a fitness disadvantage compared to the ancestor, although the difference was statistically not significant (Fig. 4B). Similar results were obtained for the permuted fluorescent combination (Fig. S1). These results demonstrated that the evolved isolates had a fitness advantage over the ancestor specifically in the plant root environment. Furthermore, the loss of fitness in a different, non-selective environment suggests an evolutionary cost of adaptation to the plant roots (Bennett and Lenski, 2007; Elena and Lenski, 2003; Van den Bergh et al., 2018).

**Fig. 4:**
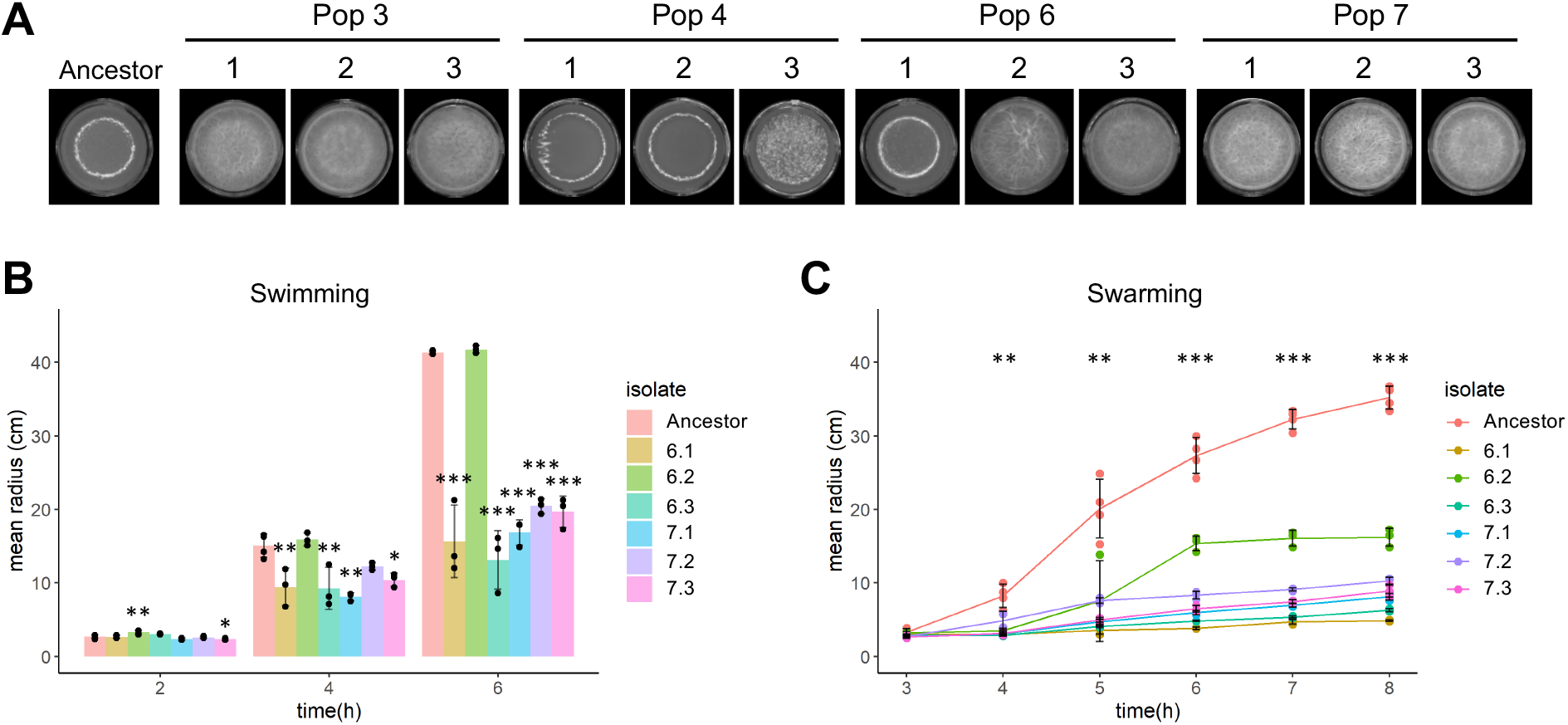
Two evolved isolates from transfer 30 outcompete the ancestor on the root but suffer a fitness disadvantage under shaking conditions in LB + xylan. Pairwise competitions between ancestor (magenta) and evolved isolates (green) during root colonization (**A**) and in LB xylan (0.5 %) under shaking conditions (B) for 48 h. In **A** and **B**, the bar plots show the starting ratio of the evolved isolate and ancestor in the mix, and the observed ratios after 48 h. Bars represent the mean (N=3-4), the error bars represent the standard deviation, and the points show the replicates. For statistical analysis, the relative fitness (r) of the evolved isolates was calculated by comparing the frequency of the evolved isolate at the beginning and at the end of the competition experiment. The log2-transformed relative fitness values were subjected to a One-sample *t*-test to test whether the mean was significantly different from 0. *: P<0.05. (C) *A. thaliana* roots colonized by a 1:1 mix of ancestor and evolved isolates imaged by CLSM. Both fluorescence combinations are shown. The top row shows the overlay of the fluorescence channels and the bright field image. Images are representative of three independent *A. thaliana* seedlings. Color codes are shown at the top. Scale bar is 50 µm.

### Evolved isolates harbor mutations in genes related to different bacterial traits

To identify the genetic changes contributing to the increased root colonization (Fig. 3) and fitness advantage over the ancestor during root colonization (Fig. 4A and C), the genomes of selected evolved isolates were re-sequenced. To represent independent populations, the isolates from populations 6 and 7 at T30 were included. Furthermore, to track molecular evolution over time, the three isolates from population 7 at T12 and T18 were also re-sequenced. Finally, one isolate from population 1 (Ev1.1) at T30 was included for re-sequencing due to its “Smooth” colony morphology and reduced root colonization (Fig. S2). In the 13 re-sequenced isolates, we observed in total 51 unique mutations of which 37 were non-synonymous (Table S1). Isolate Ev1.1 harbored several mutations in *gtaB* encoding a UTP-glucose-1-phosphate uridylyltransferase that synthesizes a nucleotide sugar precursor essential for the biosynthesis of exopolysaccharides (Varon et al., 1993) and for the synthesis of wall teichoic acids and lipoteichoic acid (Lazarevic et al., 2005). Two of the three isolates from population 6 at T30 (isolate Ev6.1 and Ev6.3) harbored a non-synonymous point mutation in the *fliM* gene, encoding a flagellar motor switch protein, part of the basal body C-ring controlling the direction of flagella rotation (Guttenplan et al., 2013). All three isolates in population 7 at T30 harbored a mutation in the intergenic region upstream of the *sinR* gene encoding a transcriptional repressor of the genes responsible for matrix production (Chu et al., 2006; Kearns et al., 2005). Interestingly, this mutation was also present in the three isolates in population 7 at T18 and in one of the isolates in this population at T12, suggesting that this mutation arose rather early in the EE and rose to a high frequency in population 7. Indeed, sequencing of the seven endpoint populations (i.e. the populations from T30) revealed that this mutation upstream from *sinR* was fixed in population 7 at the final time point, i.e. the mutation had reached a frequency of 1 in this population at T30 (Table S2). Furthermore, a nonsynonymous mutation within the *sinR* gene was detected at high frequencies in populations 2 and 3. In addition, nonsynonymous mutations in genes related to flagellar motility were besides population 6 also observed in populations 3, 4 and 5 (*fliF*, *fliK*, *fliM* and *hag*). Finally, mutations in genes related to cell wall metabolism (*gtaB*, *tagE* and *walK*, encoded functions according to SubtiWiki (Zhu and Stülke, 2018)) were identified across all seven populations (Table S2). The detection of mutations within (or upstream of) genes related to biofilm formation, motility and cell wall metabolism across independent populations supports a role of these mutations in the adaptation of *B. subtilis* to *A. thaliana* roots.

### Evolved isolates show altered pellicle biofilm formation in response to plant polysaccharides

The fitness advantage of selected evolved isolates over the ancestor during root colonization (Fig. 4A and C) and the detected mutations in the evolved isolates and endpoint populations (Table S1 and S2) confirm our initial hypothesis, that *B. subtilis* adapted to the plant root by acquiring mutations that conferred a fitness advantage over the ancestor during root colonization.

Next, we wanted to elucidate which bacterial traits were altered during such adaptation to the plant root. For this purpose, evolved isolates from the final transfer (T30) were subjected to further phenotypic characterization. Given the detected mutations (Table S1 and S2) and that both biofilm formation and motility have been shown to be important for successful root colonization by *B. subtilis* (Allard-Massicotte et al., 2016; Beauregard et al., 2013; Chen et al., 2013; Tian et al., 2021), we hypothesized that these two bacterial traits would be under selection during the adaptation to the plant roots. To this end, plant polysaccharides (PPs) including xylan have been shown to induce biofilm formation in *B. subtilis* in a non-biofilm inducing medium (Beauregard et al., 2013). One way of adapting to the plant root could thereby be through enhanced biofilm formation in response to such PPs. To test whether the improved productivity on the root by the evolved isolates was associated with more robust biofilm formation in response to PPs, the ancestor and evolved isolates were tested for pellicle biofilm formation, a biofilm formed at the medium-air interface (Branda et al., 2001), in LB supplemented with xylan (LB + xylan). Importantly, a rich medium (LB) rather than the minimal medium (MSNg) was used in this assay to provide the bacteria with plenty of nutrients, allowing us to assess only the ability of the evolved isolates to form biofilm in response to xylan, and not the ability to utilize xylan for growth. We observed that a few isolates from T30 developed a pellicle biofilm similar to the ancestor, i.e. Ev4.1, Ev4.2 and Ev6.1 (Fig. 5A). In contrast, the remaining isolates developed more robust pellicles with highly structured wrinkles indicative of enhanced matrix production. Especially the three isolates from population 7 developed hyper-robust, white pellicles, consistent with the Snow-type colony morphology observed for these isolates (Fig. 2). The biofilms developed in response to xylan by the evolved isolates generally correlated with their productivity on the root. For example, isolates Ev4.1, Ev4.2 and Ev6.1 developing similar pellicles as the ancestor and isolates Ev7.1, Ev7.2 and Ev7.3 forming hyper-wrinkled, robust pellicles in response to xylan were among the ones showing the smallest and largest increase in individual root colonization (Fig. 3 and 5A), respectively. This is in accordance with Chen *et al*. (2013) demonstrating that the ability of *B. subtilis* mutants to form robust biofilms *in vitro* correlated with that on the root. These results suggest that improved productivity on the root was associated with robust biofilm formation in response to xylan. To test whether this enhanced biofilm formation was specific to the presence of PPs, the ancestor and evolved isolates were tested for the ability to form pellicles in LB in the absence of xylan. In this medium, the pellicles developed by both the ancestor and evolved isolates were less robust (Fig. S3). For most isolates, the improved biofilm formation was specific to the presence of PPs, while the isolates from population 7 displayed robust biofilms also in the absence of plant compounds suggesting a general improvement in biofilm formation in these isolates.

**Fig. 5:**
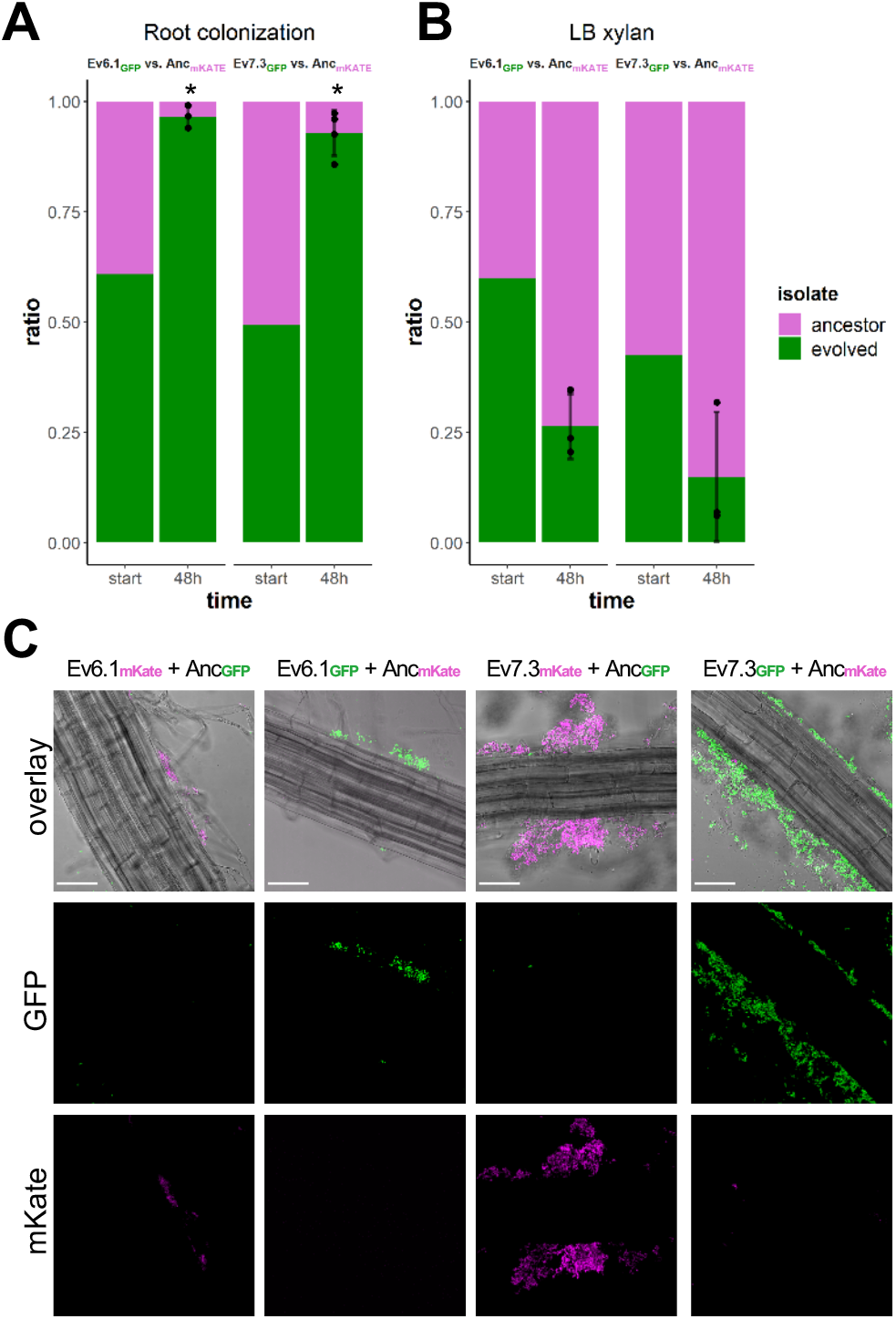
Evolved isolates from T30 show altered pellicle biofilm formation in response to plant polysaccharides and impaired motility. (**A**) Ancestor and evolved isolates from transfer 30 were inoculated into LB + 0.5 % xylan at a starting OD_600_ of 0.05 in 24-well plates. Images were acquired after 48 h incubation at 30 °C using a stereomicroscope. Each image is representative of four replicates. Evolved isolates were tested for swimming (**B**) and swarming (**C**) motility in LB medium supplemented with 0.3 or 0.7 % agar, respectively. (**B**) Bars represent the mean (N=3-4) and error bars represent standard deviation. (**C**) Lines represent the mean (N=2-4) and error bars the standard deviation. For the motility assays, the following statistical analysis applies: For each time point, an ANOVA was performed on the log10-transformed data followed by a Dunnett’s Multiple Comparison test with the ancestor as the control. For swarming motility, the asterisks show the least significance observed for the given time point. At 3 h, only isolate 7.3 was significantly reduced in swarming motility. *: P<0.05, **: P<0.01, ***: P<0.001.

### Evolved isolates show reduced swarming and swimming motility

To test whether the evolved isolates were affected in motility, the ancestor and evolved isolates from population 6 and 7 were tested for two types of motility: swimming motility, a single cell movement in aqueous environments powered by flagella rotation and swarming motility which is associated with a rapid multicellular movement of hyper-flagellated cells across a surface facilitated by self-produced surfactin (Kearns, 2010). Interestingly, most isolates were significantly impaired in both swimming and swarming motility (Fig. 5B and C). Swimming motility was observed for the ancestor and evolved isolates after 4 h (Fig. 5B). However, after 6 h only the ancestor and Ev6.2 had reached the edge of the petri dish, while the remaining isolates reached at the most half of the swimming distance of the ancestor. Swarming was observed for the ancestor after 4 h which continued until the expanding colony almost reached the edge of the petri dish after 8 h (Fig. 5C). In contrast, the evolved isolates showed reduced or a complete lack of swarming throughout the experiment. The evolution of motility-impaired isolates in independent populations could indicate that motility is not important for root colonization in the selective environment. Notably, during the EE the 48-well plates were continuously shaking at 90 rpm. We speculated, that these mildly shaking conditions could allow the bacteria to get into contact with the root by chance and thereby reducing the impact of motility on root colonization in the selective environment. To test whether motility is important during root colonization under shaking conditions, the ancestor (here referred to as “WT”) was competed against a Δ*hag* mutant, deficient in the production of the flagellin protein, for three successive rounds of root colonization under static or shaking conditions (200 rpm). Under static conditions, the Δ*hag* mutant was significantly outcompeted by the WT (Fig. S4). In contrast, under shaking conditions, the Δ*hag* mutant was able to co-colonize the root to similar levels as the WT. These results demonstrate that motility is important for competition on the root under static conditions but is not required under shaking conditions. Thereby, impaired motility of several of the evolved isolates is not expected to negatively influence the fitness of these isolates in the selective environment.

### Evolved isolates show distinct growth profiles in a plant-mimicking environment

The minimal medium (MSNg) used in the EE should render the bacteria dependent on root exudates and dead plant material to survive. A simple way of adapting to this selective environment could be through the enhanced utilization of available plant compounds. To test this, we used a modified version of the minimal medium employed during the EE. Instead of 0.05 % glycerol, the MSN medium was supplemented with 0.5 % cellobiose (MSNc). Cellobiose is a disaccharide and a product of partial hydrolysis of cellulose, found in plant cell walls (Beauregard et al., 2013; Endler and Persson, 2011). In addition, MSNc was supplemented with 0.5 % xylan. The ancestor showed a growth profile typical of bacterial growth under planktonic conditions (Fig. S5). In contrast, several evolved isolates displayed distinct growth profiles, including 3.2, 7.1, 7.2 and 7.3, which showed no decline phase, but instead displayed a prolonged stationary phase. When analyzing the carrying capacity (K), several isolates showed significantly increased carrying capacity (all three isolates from populations 3 and 7), while few isolates showed significantly decreased carrying capacity (Ev4.2, Ev4.3 and Ev6.2). While cellobiose and xylan do not completely represent the plant compounds present in the selective environment, these results suggest that adaptation to the plant root could also be facilitated through the altered utilization of plant compounds.

### An evolved isolate shows increased colonization of *A. thaliana* roots in the presence of a synthetic, soil-derived community

During the EE, *B. subtilis* was adapted to the plant root alone – in the absence of other microbes. This selective environment is far from its natural habitat in the rhizosphere, where *B. subtilis* encounters other microbial residents. In fact, the ancestor DK1042 is a derivate of the wild strain NCIB 3610, originally isolated from hay infusion (Cohn, 1930; Zeigler et al., 2008). To this end, we wondered how the pro-longed adaptation of *B. subtilis* to the plant root environment in the absence of other microbial species affected the ability to colonize the root in the presence of soil microbes. The ancestor and Ev7.3 were tested for their ability to colonize *A. thaliana* roots in the presence of a synthetic, soil-derived community (Lozano-Andrade et al., 2021). This community comprises four bacterial species, *Pedobacter* sp., *Rhodococcus globerulus*, *Stenotrophomas indicatrix* and *Chryseobacterium* sp. which were previously isolated from soil samples that also contained *B. subtilis*, thereby representing bacterial soil inhabitants that *B. subtilis* would normally encounter in nature. The isolate Ev7.3 was chosen for this test since it was highly adapted to the selective environment, i.e. the isolate displayed significantly increased individual root colonization (Fig. 3) and outcompeted the ancestor during competition on the root, where it formed a robust biofilm along the root (Fig. 4A, C). To capture any potential difference in the establishment on the root, here defined as root colonization after 48 h, between *B. subtilis* ancestor and Ev7.3 in the presence of the community, the ancestor or Ev7.3 was co-inoculated with the community in four different ratios: 0.1:1, 1:1, 10.1 and 100:1 of *B. subtilis* and community, respectively. When *B. subtilis* was initially under-represented or highly in excess, i.e. inoculation ratio 0.1:1 and 100:1, respectively, no significant difference was observed in the establishment on the root between *B. subtilis* ancestor and Ev7.3 within the same inoculation ratio (Fig. 6). In contrast, when co-inoculated with the community in intermediate ratios, i.e. 1:1 and 10:1, isolate Ev7.3 showed significantly enhanced establishment on the root compared to the ancestor. Since Ev7.3 displayed increased carrying capacity in MSNc + xylan in monoculture compared to the ancestor (Fig. S5), we wondered whether the enhanced establishment on the root by Ev7.3 in the presence of the community could be partly attributed to improved utilization of plant compounds. Indeed, growth profiles in MSNc + xylan of the ancestor or Ev7.3 in co-culture with the community revealed that Ev7.3 displayed a significantly increased carrying capacity at inoculation ratio 1:1, 10:1 and 100:1 compared to the ancestor (Fig. S6). Finally, *in vitro* confrontation assays on LB agar (1.5 %) showed no major difference in the inhibition of the community members by Ev7.3 compared to the ancestor (Fig. S7). Taken together, these results show that even though *B. subtilis* was adapted to the plant root alone, isolate Ev7.3 displayed increased root colonization also in the presence of a synthetic, soil-derived community under certain inoculation ratios, possibly mediated by robust biofilm formation on the root and enhanced utilization of plant compounds.

**Fig. 6:**
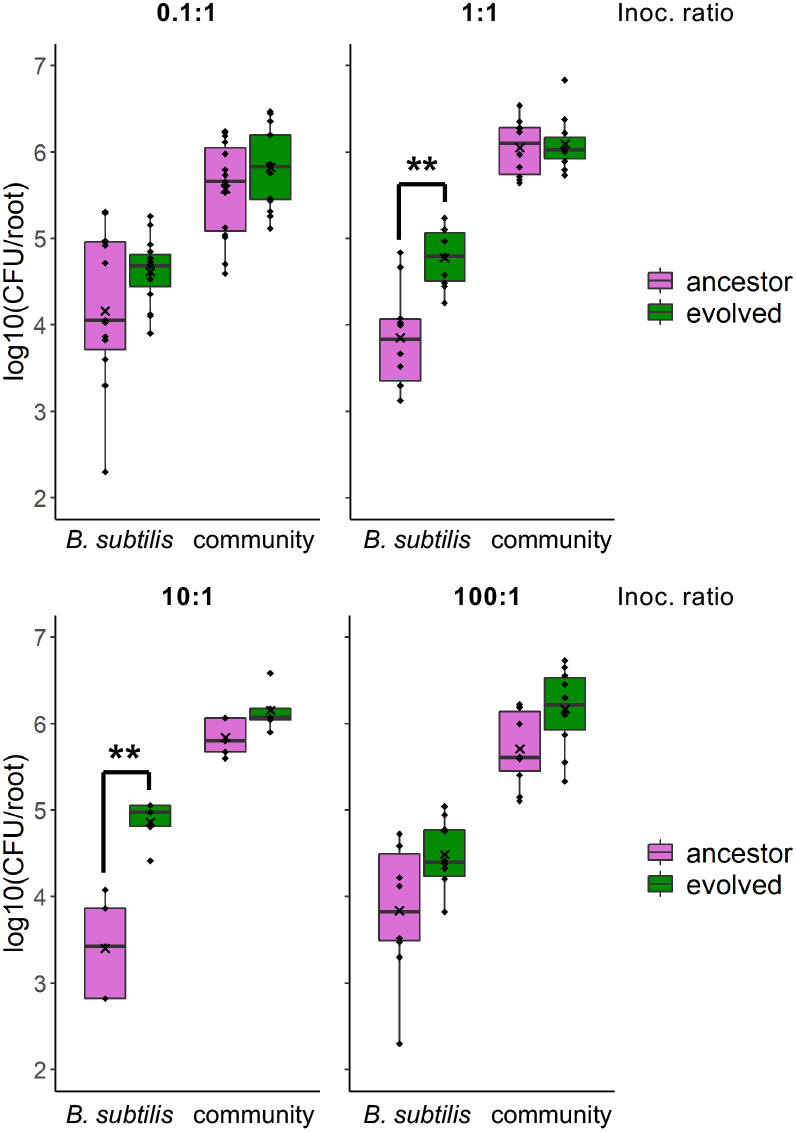
Root colonization by *B. subtilis* ancestor and isolate Ev7.3 in the presence of a synthetic, soil-derived community. The ancestor and evolved isolate, Ev7.3, were tested for the ability to colonize the root in the presence of a synthetic, soil-derived bacterial community. *B. subtilis* ancestor or Ev7.3 and the community were co-inoculated onto *A. thaliana* roots in four different ratios: 0.1:1, 1:1, 10:1 and 100:1 of *B. subtilis* and community, respectively. Root colonization after 48 h was quantified as log10-transformed productivity (CFU/root). Each plot shows the resulting root colonization at the given inoculation ratio of *B. subtilis* (left) and the co-inoculated community (right). Magenta: Ancestor and the corresponding community co-inoculated with the ancestor. Green: Ev7.3 and the community co-inoculated with Ev7.3. The cross represents the mean (N=5-15). Within each inoculation ratio, statistical significance between *B. subtilis* ancestor and Ev7.3, and between the communities co-inoculated with the ancestor or with Ev7.3 was tested with a Two-sample *t*-test (Welch’s Two-sample *t*-test when unequal variance). **: P<0.01.

## Discussion

Several studies have reported experimental evolution as a powerful tool to explore how bacteria adapt to ecologically relevant environments. A recent study investigated the adaptive response of the PGPR *Pseudomonas protegens* to the *A. thaliana* rhizosphere in a sand system, which revealed mutations in genes encoding global regulators and genes related to motility and cell surface structure across independent populations (Li et al., 2021b) and during such adaptation, the initially plant-antagonistic *P. protegens* bacterium evolved into mutualists (Li et al., 2021a). Furthermore, Lin *et al*. (2021) observed that adaptation of *Bacillus thuringiensis* to *A. thaliana* roots under hydroponic conditions led to the evolution of multicellular aggregating phenotypes, which, surprisingly, in certain lineages were accompanied by enhanced virulence against the *Galleria mellonella* larvae. Here, we employed experimental evolution to study the adaptation of *B. subtilis* to *A. thaliana* roots under hydroponic conditions. Our initial hypothesis was that *B. subtilis* would adapt to the plant roots by acquiring mutations that would provide the bacteria with a fitness advantage over the ancestor during root colonization. We could demonstrate that *B. subtilis* rapidly adapted to the plant roots, as observed by evolved isolates displaying improved root colonization relative to the ancestor already after 12 transfers and the detection of genetic changes in evolved isolates from transfer 12, 18 and 30. In addition, competition between the ancestor and two selected evolved isolates from the final transfer (T30) on the root revealed that both evolved isolates had a fitness advantage over the ancestor during root colonization, thereby confirming our hypothesis.

Further phenotypic characterization of the evolved isolates from the final transfer revealed that most isolates across independent populations developed more robust biofilms in response to the plant polysaccharide xylan compared to the ancestor. Except for isolate Ev3.3, the robust biofilm formers tended to be increased in individual root colonization, indicating that robust biofilm formation is associated with adaptation to the plant root. Motility represents an important trait for many bacteria as it allows them to explore the environment for nutrients and escape unfavorable conditions. Of relevance to the adaptation of *B. subtilis* to plant roots, motility has been shown to be important for root colonization of different plant species under different conditions. For example, a *B. subtilis* Δ*hag* mutant was shown to be delayed or reduced in *A. thaliana* root colonization under hydroponic conditions as well as in tomato root colonization under vermiculites pot conditions (Allard-Massicotte et al., 2016; Tian et al., 2021). Yet, we found that five out of six isolates from two independent populations were impaired in both swimming and swarming motility, indicating that motility is not important for root colonization in the selective environment of the EE, i.e. under hydroponic, shaking conditions. Indeed, this was verified in a competition experiment between a non-motile Δ*hag* mutant and the WT, revealing that motility is not required for root colonization under shaking conditions. In contrast to our observations, Li *et al*. (2021) observed several evolved isolates of *P. protegens* improved in swimming motility following adaptation to the *A. thaliana* rhizosphere in a sand system (Li et al., 2021b), supporting that in sand (and thus possibly also in soil), motility is indeed important for root colonization and is therefore selected for.

In *B. subtilis*, motility and biofilm formation are incompatible processes: *B. subtilis* can exist as single, motile cells or in chains of sessile, matrix-producing cells which is regulated by an epigenetic switch involving SinR (Chai et al., 2010; Vlamakis et al., 2008). The enhanced biofilm formation and impaired motility of isolates from populations 6 and 7 (Ev6.3, Ev7.1, Ev7.2 and Ev7.3) could thereby indicate a possible biofilm-motility trade-off. An inverse evolutionary trade-off between biofilm formation and motility was observed when the opportunistic pathogen *Pseudomonas aeruginosa* was subjected to repeated rounds of swarming which lead to the evolution of hyper-swarmers that were impaired in biofilm formation (van Ditmarsch et al., 2013). Considering that the ability to form robust biofilm *in vitro* was shown to positively correlate with root colonization in *B. subtilis* (Chen et al., 2013), and the demonstration that motility is not important for root colonization under shaking conditions, a possible biofilm-motility trade-off could provide *B. subtilis* with enhanced fitness during root colonization in the selective environment. Indeed, isolate Ev7.3, which developed hyper-robust biofilms in LB + xylan and was impaired in motility, significantly outcompeted the ancestor during root colonization.

Re-sequencing of selected evolved isolates revealed that Ev7.1, Ev7.2 and Ev7.3 (from transfer 30) all harbored a single nucleotide polymorphism (SNP) two base pairs upstream from the start codon of the *sinR* gene (Agarwala et al., 2018), encoding a transcriptional repressor of matrix genes (Chu et al., 2006; Kearns et al., 2005). This SNP is located in the spacer region between the Shine Dalgarno sequence and the start codon in the ribosome binding site. Interestingly, the nucleotide composition of the spacer sequence has been shown to influence translation efficiency (Liebeton et al., 2014). The SNP upstream from *sinR* might thereby potentially affect the translation efficiency from the mRNA transcript, resulting in reduced levels of SinR. Reduced levels of SinR could in turn result in increased expression of matrix genes. This is supported by Richter *et al*. (2018) who demonstrated that a Δ*sinR* mutant shows increased matrix gene expression, and by Subramaniam *et al*. (2013) reporting that SinR translation and therefore protein level affects matrix gene expression. Potential increased matrix production caused by this mutation could contribute to the Snow-type colony morphology, as this colony morphology was exclusively observed for isolates harboring this mutation. Furthermore, in accordance to the robust biofilm formation and increased root colonization observed for Ev7.1, Ev7.2 and Ev7.3, a Δ*sinR* mutant was shown to form a hyper-robust biofilm in biofilm-inducing medium as well as on tomato roots (Chen et al., 2013), supporting the possible relevance of this mutation for the observed phenotypes of these isolates. Based on these previous studies, we therefore speculate that the mutation upstream from *sinR* results in increased matrix gene expression, which in turn enables more robust biofilm formation and increased root colonization as observed for the three isolates in population 7. These three isolates did not harbor mutations in motility-related genes. However, besides a possible effect of reduced SinR levels on the epigenetic switch (Chai et al., 2010) that could lock the cells in a sessile, matrix-producing stage, a potential reduction in SinR levels leading to overexpression of the *eps* operon could possibly reduce motility due to the EpsE clutch (Blair et al., 2008). Such mutation and the observed corresponding phenotypes could be responsible for the biofilm-motility trade-off, and be an example of antagonistic pleiotropy (Elena and Lenski, 2003) in which the same mutation is beneficial in one environment, i.e. during root colonization under shaking conditions, but disadvantageous in another, i.e. where motility is required for survival. In addition, this mutation affecting a biofilm regulator could possibly explain why Ev7.1, Ev7.2 and Ev7.3 show improved biofilm formation also in the absence of xylan.

Isolate Ev6.1 and Ev6.3 harbored a non-synonymous point mutation in the *fliM* gene, which was not present in Ev6.2. This gene encodes a flagellar motor switch protein, part of the basal body C-ring controlling the direction of flagella rotation (Guttenplan et al., 2013). Interestingly, Ev6.1 and Ev6.3 were impaired in both forms of motility, while Ev6.2 showed similar swimming as the ancestor and was less affected in swarming. We speculate that the R326I substitution affects the function of FliM and consequently the flagellar machinery, resulting in hampered motility in these two isolates. Since we showed that motility was not important for root colonization under shaking conditions, a mutation hampering motility could provide the bacterium a fitness advantage during the adaptation to *A. thaliana* roots due to the reduced cost of this apparently redundant trait. However, we do not expect the mutation in *fliM* to result in reduced cost; it merely changes an amino acid in a protein part of the flagellar machinery. Other mutations in the population 6 isolates must explain the robust biofilm formation by Ev6.2 and Ev6.3 and the fitness advantage of Ev6.1 over the ancestor during root colonization. For example, isolate Ev6.1 and Ev6.3 harbor a mutation in *kinA* encoding a two-component sensor kinase which once activated initiates the phosphorelay leading to phosphorylation of the master regulator Spo0A (Jiang et al., 2000).

The isolate from population 1 at transfer 30 (Ev1.1) harbored a frameshift mutation in the *rsiX* locus, encoding an anti-sigma factor controlling the activity of SigX (Zhu and Stülke, 2018). Inconsistent with the smooth morphology and reduced root colonization of this isolate, an Δ*rsiX* mutant was shown to have increased *eps* expression (Martin et al., 2020). However, Ev1.1 additionally harbored several mutations in *gtaB* encoding a UTP-glucose-1-phosphate uridylyltransferase involved in the biosynthesis of a nucleotide sugar precursor for EPS biosynthesis (Varon et al., 1993). A study conducted by Reverdy *et al*. (2018) showed that acetylation of GtaB is important for biofilm formation of *B. subtilis* and that a *gtaB* mutant was reduced in pellicle formation. In addition, Xu *et al*. (2019) showed that a Δ*gtaB* mutant of *B. velezensis* SQR9 was significantly decreased in colonization of cucumber roots compared to the WT, although the effect of *gtaB* on root colonization may be species dependent. We therefore speculate, that Ev1.1 may have first gained the mutation in *rsiX* and was selected for due to increased *eps* expression, while later on the increased matrix production was reverted by the mutations in *gtaB*, resulting in a non-functional protein and thereby reduced precursors for EPS production, which was selected for due to reduced cost. In accordance with the mutations observed in this study, a non-synonymous mutation in *sinR* in two isolates forming Snow-type colonies and increased in root colonization as well as several mutations in *gtaB* in two isolates with a Smooth colony morphology were observed in our recent study on diversification of *B. subtilis* during adaptation to *A. thaliana* roots (Blake et al., 2021b).

To get a more general insight into the mutations arising in *B. subtilis* during EE on *A. thaliana* roots, the seven endpoint populations were also sequenced. This revealed mutations within (or upstream of) genes related to biofilm formation (*sinR*), flagellar motility (*fliF*, *fliK*, *fliM* and *hag*) and cell wall metabolism (*gtaB*, *tagE* and *walK*) (Zhu and Stülke, 2018) across independent populations. Taken together, our findings of evolved isolates displaying altered biofilm formation and motility properties and the detection of mutations within (or upstream of) genes related to biofilm formation and motility in single evolved isolates as well as across independent endpoint populations indicates that adaptation of *B. subtilis* to *A. thaliana* roots under the employed conditions is associated with alterations in these two bacterial traits. While we found that the phenotypic and genetic changes of Ev6.1 and Ev7.3 conferred a fitness advantage over the ancestor during root colonization, adaptation to one certain environment may be accompanied by a loss of fitness in other environments (Elena and Lenski, 2003). This has been demonstrated for *Escherichia coli* which following adaptation to low temperature showed reduced fitness at high temperature (Bennett and Lenski, 2007). In the example of the evolution of hyperswarmers of *P. aeruginosa*, the hyperswarmer clones outcompeted the ancestor in swarming, but lost in biofilm competitions (van Ditmarsch et al., 2013). In this study, we demonstrate that adaptation of *B. subtilis* to *A. thaliana* roots is accompanied by an evolutionary cost. When Ev6.1 and Ev7.3 each were competed against the ancestor in LB + xylan under shaking conditions, i.e. an environment where plant compounds are present but biofilm formation is not required for survival, both evolved isolates suffered a fitness disadvantage. The observation that two evolved isolates, from independent populations and with different phenotypes and genetic changes, both suffered a fitness disadvantage in a non-selective environment might suggest the generality of such an evolutionary cost accompanying adaptation to *A. thaliana* roots.

In our EE approach, *B. subtilis* was adapted to plant roots in the absence of other microbes. In the rhizosphere environment under natural conditions, *B. subtilis* is far from being the sole microbial inhabitant. Instead, it engages in cooperative and competitive interactions with other members of the rhizosphere microbiome (Hassani et al., 2018; Kiesewalter et al., 2021). We tested whether the evolved isolate, Ev7.3, displaying increased root colonization in the selective environment relative to the ancestor, would also show improved establishment on the root under more ecologically complex conditions. We found that in the presence of a synthetic, soil-derived community, Ev7.3 displayed enhanced establishment on the root compared to the ancestor in two out of four inoculation ratios. This enhanced establishment on the root by Ev7.3 is not expected to be caused by altered antagonistic activities towards the community members. First, no major changes in the inhibition of the community members were observed in confrontation colony assays. Secondly, an increased number of Ev7.3 cells on the root did not cause a reduction in the co-colonizing community. Finally, Ev7.3 did not harbor mutations in genes directly related to secondary metabolite production. Instead, enhanced establishment on the root by Ev7.3 in the presence of the community is possibly enabled by robust biofilm formation facilitating stronger attachment to the root and enhanced utilization of plant compounds. Interestingly, a study by Molina-Santiago *et al*. (2019) showed that compared to a Δmatrix mutant, co-inoculation of *B. subtilis* WT with *Pseudomonas chlororaphis* on melon leaves enabled co-localization of the two species as well as closer attachment of *B. subtilis* to the leave surface (Molina-Santiago et al., 2019). The robust biofilm formed on the root by Ev7.3 possibly facilitated by increased matrix production may thereby not exclude the community members on the root but could rather allow them to incorporate into the matrix. This could also explain why the enhanced establishment of *B. subtilis* Ev7.3 on the root did not cause a reduction in the number of community cells attached to the root. Alternatively, the community may not be majorly affected by any difference in establishment on the root between the ancestor and Ev7.3 due to the low abundance of *B. subtilis* relative to the community. Further work is needed to elucidate the interactions between *B. subtilis* and this synthetic community during root colonization. In summary, these findings suggest that even though *B. subtilis* was evolved on *A. thaliana* in the absence of other microbes, it became highly adapted to the plant root environment enabling better establishment on the root also when the ecological complexity increases. How genetic adaptation to the plant root in the absence of other microbial species differs from adaptation to plant root environments with varying levels of ecological complexity is the scope of future studies.

The formation of root-associated biofilms is important for the biocontrol efficacy of *B. subtilis* (Chen et al., 2013). From an applied perspective, experimental evolution of *B. subtilis* on plant roots represents a novel approach for developing strains with enhanced root attachment capacities for agricultural use. However, a biofilm-motility tradeoff as observed here may be undesirable when developing biocontrol agents due to the growing evidence of motility as an important trait for bacterial root colonization in soil systems (Li et al., 2021b; Tian et al., 2021). The phenotypes associated with adaptation of *B. subtilis* to *A. thaliana* roots presented here as well as the accompanying evolutionary cost and the increased root colonization also in the presence of resident soil bacteria highlight the importance of considering the selective environment if evolving PGPR for biocontrol purposes.

## Limitations of the Study

This study on evolutionary adaptation of *B. subtilis* to *A. thaliana* roots under hydroponic conditions revealed that *B. subtilis* rapidly adapted to the plant root environment as observed by improved root colonizers already after 12 transfers. Moreover, we found that one selected evolved isolate displayed increased root colonization also in the presence of resident soil bacteria. The findings from this study thereby highlight experimental evolution as an approach to develop *B. subtilis* strains with improved root colonization capacities to potentially support a sustainable agricultural production. However, such plant root adaptation might be condition-specific, and we do not know whether the evolved isolates also display increased root colonization under soil conditions reminiscent of those the bacteria will encounter under greenhouse or field conditions. To this end, motility has been shown to be important for root colonization under diverse conditions, including soil and sand conditions (Gao et al., 2016; Tian et al., 2021). Thereby, the biofilm-motility trade-off observed for several of the evolved isolates might be undesirable in a biocontrol strain used under field or greenhouse conditions. Future studies will therefore test the evolved isolates for root colonization under greenhouse soil conditions to reveal whether adaptation to plant roots under the simple axenic, hydroponic conditions employed in this study, manifests in increased root colonization also under agriculturally relevant conditions.

## Authors’ contributions

M.N.C. and Á.T.K. conceived the project; M.N.C. and C.B. performed experiments; M.N.C. and M.L.S analyzed the data; G.M. performed the re-sequencing of single isolates and the belonging data analysis; G. H. and Y. W. performed re-sequencing of endpoint populations and the belonging data analysis; M.N.C. and Á.T.K. wrote the manuscript with feedback from all authors.

## Acknowledgements

We thank Carlos N. Lozano-Andrade for the four bacterial species constituting the synthetic community. The work was supported by a DTU Bioengineering start-up fund to ÁTK. Fundings from Novo Nordisk Foundation for the infrastructure “Imaging microbial language in biocontrol (IMLiB)” (NNFOC0055625) and within the INTERACT project of the Collaborative Crop Resiliency Program (NNF19SA0059360) are acknowledged. The position of M.L.S. is financed by the Danish National Research Foundation (DNRF137) for the Center for Microbial Secondary Metabolites. G.H. and Y.W. were supported by China National GeneBank (CNGB).

## Supplemental item titles

**Table S1:** The table shows detected mutations in the re-sequenced genomes of evolved isolates. The functions of the gene products were retrieved from the SubtiWiki Database (Zhu and Stülke, 2018). T = Transfer.

**Table S2:** The table shows detected mutations in the sequenced endpoint populations. The functions of the gene products were retrieved from the SubtiWiki Database (Zhu and Stülke, 2018). T = Transfer.

## Methods

### Data and code availability

The sequencing data for single evolved isolates has been deposited into the NCBI Sequence Read Archive (SRA) database under BioProject accession number: PRJNA705352, and sequencing data for endpoint populations into CNGB Sequence Archive (CNSA) (Guo et al., 2020) of China National GeneBank DataBase (CNGBdb) (Chen et al., 2020) with accession number CNP0002416.

Other data reported in this study is available from the lead contact upon reasonable request. This study does not report any new, original code.

## EXPERIMENTAL MODELS AND SUBJECT DETAILS

### Bacterial strains and culture media

*Bacillus subtilis* DK1042 strain, an easily transformable derivative of the undomesticated *B. subtilis* NCBI 3610 (Konkol et al., 2013), was used as ancestor for the experimental evolution. For pairwise competitions between the ancestor and evolved isolates, and for root co-colonization with a synthetic, soil-derived community, TB500.1 and TB501.1, that were previously created by transforming DK1042 with pTB497.1 and pTB498.1 (Dragoš et al., 2018a; Mhatre et al., 2017), respectively, were used as the ancestor. In addition, selected evolved isolates derived from the ancestor were transformed with plasmids pTB497.1 and pTB498.1. These plasmids harbor the *gfp* and *mKATE* gene, respectively, under the control of the hyper-spank (constitutive) promoter and a spectinomycin resistance gene within the flanking regions of the *amyE* gene. Transformants were identified by selecting for spectinomycin resistance and double crossovers were verified by the loss of amylase activity.

The four bacterial species, *Pedobacter* sp.*, Rhodococcus globerulus*, *Stenotrophomas indicatrix* and *Chryseobacterium* sp., constituting a synthetic, soil-derived community were previously acquired in Dyrehaven, Kongens Lyngby, Denmark (55° 47’ 19.68” N, 12° 33’ 29.88” E), as described (Lozano-Andrade et al., 2021). An overview of the strains used in this study is included in the Key Resources Table. *B. subtilis* strains were routinely grown overnight (ON) in Lysogeny Broth (LB; LB-Lennox, Carl Roth, Germany; 10 g/L tryptone, 5 g/L yeast extract and 5 g/L NaCl) at 37 °C while shaking at 220 rpm. When relevant, spectinomycin was added at a final concentration of 100 µg/mL. The bacterial species used for the synthetic, soil-derived community were grown for 48 h in 0.1 % (w/v) Tryptic Soy Broth (TSB; Sigma-Aldrich, St. Louis, Missouri, USA) at room temperature. For the experimental evolution on plant roots and root colonization assays, a minimal salts nitrogen glycerol (MSNg) medium was used. MSNg was prepared as follows: The base was prepared by adding 0.026 g KH_2_PO_4_, 0.061 g K_2_HPO_4_, 2.09 g MOPS and 0.04 g MgCl_2_ x 6H_2_O per 100 mL dH_2_O and adjusting the pH to 7.0 using KOH. The base was autoclaved, cooled down to room temperature and then supplemented with 0.05 mL of 100 mM MnCl_2_, 0.1 mL of 1 mM ZnCl_2_, 0.1 mL of 2 mM thiamine, 0.1 mL of 0.7 M CaCl_2_, NH_4_Cl_2_ to a final 0.2 % and glycerol to a final 0.05 %. Growth of *B. subtilis* ancestor and evolved isolates alone or in co-culture with the synthetic, soil-derived community was monitored in MSNc + xylan, which was prepared similarly to MSNg except that instead of glycerol, cellobiose and xylan were added to a final concentration of 0.5 %. Pellicle biofilm formation was assessed in LB medium with or without a supplementation of xylan (0.5 %) (w/v). Pairwise competitions between the ancestor and evolved isolates were evaluated in LB + xylan (0.5 %).

### Plant material

*Arabidopsis thaliana* Col-0 seedings were used as plant host for the experimental evolution and root colonization assays. The seeds were surface sterilized in a 2% (v/v) sodium hypochlorite solution (VWR, Radnor, Pennsylvania, USA) and shaken on an orbital mixer for 12 minutes. The seeds were washed five times in sterile, distilled water. Approximately 15 sterilized seeds were carefully pipetted onto pre-dried MS agar plates (Murashige and Skoog basal salts mixture, Sigma-Aldrich) (2.2 g l^-1^, pH = 5.6-5.8 supplemented with 1 % agar). Plates were sealed with parafilm and stratified at 4 °C for 3 days, and were then placed at an angle of 65 °C in a plant chamber (cycles of 16 h light at 24 °C, and 8 h dark at 20 °C). The seedlings were grown in the plant chamber for 6-8 days before use.

## METHOD DETAILS

### Experimental evolution of *B. subtilis* on *A. thaliana* seedlings

To study the evolutionary adaptation of *B. subtilis* to *A. thaliana* roots, we employed an experimental evolution (EE) setup similar to the one established by Lin et al. (2021). This setup was inspired by a long-term EE on polystyrene beads (Poltak and Cooper, 2011), but instead of beads, *A. thaliana* roots were used for successive root colonization by *B. subtilis*. The EE on plant seedlings was carried out under axenic, hydroponic conditions optimized for *B. subtilis* (Beauregard et al., 2013; Dragoš et al., 2018a; Gallegos-Monterrosa et al., 2016; Nordgaard et al., 2021; Thérien et al., 2020). Seven parallel populations were initiated by inoculating *A. thaliana* seedlings, placed in 300 µL MSNg medium in 48-well plates, with *B. subtilis* DK1042 at a starting OD_600_ of 0.02. In addition, a replicate with a sterile root in MSNg without bacterial inoculation was included as a control. The 48-well plate was incubated in a plant chamber (cycles of 16 h light at 24 °C/8 h dark at 20 °C) at mild agitation (90 rpm). After 48 h, the colonized seedling was washed twice in MSNg and transferred to fresh medium containing a new, sterile seedling enabling re-colonization. Following this approach, the seven parallel populations were serially transferred every 48 h for a total of 32 transfers. At different time points during the ongoing EE, the old seedlings were washed twice in MSNg and vortexed with glass beads to disperse the biofilm. The resulting cell suspension was used for plating to follow the productivity of the evolving populations, i.e. colony-forming unit (CFU) per root, and also preserved as frozen (−80°C) stocks for later analysis.

### Isolation of single evolved isolates and colony morphology assay

Frozen stocks of population 3, 4, 6 and 7 from three different time points during the EE (transfer 12, 18 and 30) were streaked on LB plates to obtain single colonies of evolved isolates. Transfer 30 (T30) was designated as the final transfer for the isolation of evolved isolates. From each population and time point, three colonies were randomly selected, prepared as ON cultures and saved as frozen stocks. Colony morphologies of evolved isolates were examined by spotting 2 µL ON cultures on LB agar (1.5 %) and incubated at 30 °C for 48 h. Importantly, by growing the ancestor and evolved isolates from frozen stocks in LB ON, all isolates should be in a similar physiological state, so a potential difference in colony morphology should be attributed to genetic variation.

### Root colonization assay

The evolved isolates were tested for root colonization under the same conditions applied during the EE. For individual root colonization, sterile *A. thaliana* seedlings in MSNg medium were inoculated with the ancestor or evolved isolates at a starting OD_600_ of 0.02. For pairwise competition experiments, sterile *A. thaliana* seedlings were inoculated with a 1:1 mix of the ancestor and evolved isolates with opposite, fluorescent labels. Importantly, to obtain a similar starting cell number for the competition experiment, ON cultures of the ancestor and isolate Ev6.1 were adjusted to an OD_600_ of 0.2, while Ev7.3 was adjusted to an OD_600_ of 1.0 corresponding to comparable total cell counts. For plant co-colonization by *B. subtilis* in the presence of a soil-derived synthetic community, cell suspensions of *Pedobacter* sp. D749 and *Rhodococcus globerulus* D757 were adjusted to an OD_600_ of 2.0, *Stenotrophomas indicatrix* D763 to an OD_600_ of 0.05 and C*hryseobacterium* sp. D764 to an OD_600_ of 0.1 and mixed by equal volumes (hereafter referred to as community). This mix was further adjusted to an OD_600_ of 0.2. OD-adjusted *B. subtilis* ancestor (OD_600_ of 0.2) or Ev7.3 (OD_600_ of 1.0) and community were co-inoculated in four different ratios (0.1:1, 1:1, 10:1, 100:1 of *B. subtilis* and community, respectively) into MSNg medium containing a sterile *A. thaliana* seedling. The 48-well plates of all root colonization assays were incubated in the plant chamber at mild agitation (90 rpm). After 48 h, the colonized seedling was washed twice in MSNg and either vortexed with glass beads, and the resulting cell suspension plated for CFU quantification, or transferred to a glass slide for imaging using confocal laser scanning microscopy (CLSM).

For root colonization competition between *B. subtilis* WT and the Δ*hag* mutant, ON cultures of the WT and Δ*hag* mutant with opposite fluorescent labels were mixed 1:1 and inoculated onto sterile *A. thaliana* seedlings in MSNg medium at a starting OD_600_ of 0.02. Plates were incubated in the plant chamber under static or shaking conditions (200 rpm) for 48 h. The seedlings were then washed in MSNg and transferred to fresh medium containing a new, sterile seedling and incubated under the same conditions as before (i.e. static or shaking conditions). This step was repeated once more, after which the third root was washed and vortexed with glass beads, and the resulting cell suspension was plated for CFU quantification.

### Pairwise competition experiments in LB + xylan

ON cultures of the ancestor and selected evolved isolates were adjusted to an OD_600_ of 5.0. Ancestor was mixed 1:1 and 1:5 by volume with Ev6.1 and Ev7.3, respectively, to obtain a comparable number of starting cells. 80 µL of the mix was inoculated into 20 mL LB + xylan (0.5 %) medium in 100 mL bottles, giving a starting OD_600_ of 0.02. Bottles were incubated at 37 °C while shaking at 220 rpm for 48 h followed by plating for CFU quantification.

### Biofilm formation in response to plant polysaccharides

To test for pellicle biofilm formation in response to plant polysaccharides, the ancestor or the evolved isolates were inoculated into 1.5 mL LB + xylan (0.5 %) medium in 24-well plates to a starting OD_600_ of 0.05 and incubated under static conditions at 30 °C for 48 h. In addition, the ancestor and evolved isolates were evaluated for biofilm formation in LB without supplementation.

### Motility assays

Swimming motility was tested using soft agar plates (15 mL LB with 0.3 % agar) dried for 5 minutes while swarming motility was evaluated on semi-soft agar plates (15 mL LB with 0.7 % agar) dried for 20 minutes. 2 µL ON cultures of the ancestor or evolved isolates adjusted to an OD_600_ of 0.5 were spotted in the middle of a petri dish and incubated at 37 °C. Multiple stacking was avoided in order to keep a similar humidity across all plates. Swimming and swarming motility were followed for 6 and 8 h, respectively. For each plate, motility was quantified as the averaged radius measured in four different directions.

### Growth in the presence of xylan

To monitor the growth of ancestor and evolved isolates in the presence of plant compounds, two independent ON cultures of each isolate were independently inoculated into a 96-well plate containing 100 µL MSNc + xylan (0.5 %) medium at a starting OD_600_ of 0.1. OD_600_ was monitored in a plate reader (BioTek Synergy HTX Multi-Mode Microplate Reader) every 15 min for 55 h at 24 °C under continuous shaking (orbital). To test for growth in co-culture with the community, two to three independent ON cultures of constitutively GFP-labelled *B. subtilis* ancestor and Ev7.3 were adjusted to an OD_600_ of 0.1 in MSNc + xylan (0.5 %). Cell suspensions of the four community members were adjusted in MSNc + xylan (0.5 %) to the same OD_600_ values used for the root colonization assay (see above), and the mixed community was adjusted to a final OD_600_ of 0.1. Ancestor or Ev7.3 was co-inoculated with the community in the same ratios as for the root colonization assay (0.1:1, 1:1, 10:1 and 100:1 of *B. subtilis* and community, respectively). OD_600_ and GFP were monitored in the plate reader every 15 minutes for 35 h at 24 °C under continuous shaking (orbital). In both growth assays, each well was measured at 9 (OD_600_) or 5 (GFP_485/20nm_) different points to avoid artifacts due to aggregation.

### Pairwise interactions of *B. subtilis* with community members

To study potential altered interactions with any of the four bacterial species of the community, 2 µL of ON cultures of *B. subtilis* ancestor or Ev7.3 adjusted to an OD_600_ of 0.5 was spotted on LB agar (1.5 %). On the same plate, 2 µL of cell suspensions of *Pedobacter* sp. (OD_600_ of 2.0), *Rhodococcus globerulus* (OD_600_ of 2.0), *Stenotrophomas indicatrix* (OD_600_ of 0.1) or *Chryseobacterium* sp. (OD_600_ of 0.1) were spotted at a 0.7 cm distance from the *B. subtilis* inoculum. The plates were incubated at 30 °C.

### Microscopy/confocal laser scanning microscopy (CLSM)

Bright-field images of colonies, whole pellicle biofilms and pairwise interactions were acquired with an Axio Zoom V16 stereomicroscope (Carl Zeiss, Germany) equipped with a Zeiss CL 9000 LED light source and an AxioCam MRm monochrome camera (Carl Zeiss, Germany). Images of colonized seedlings were acquired using CLSM (Leica Microsystems Confocal Microscope SP8, Germany). The seedlings were washed twice in MSNg and placed onto glass slides. Images were obtained using the 63× / 1.4 OIL objective. Fluorescent reporter excitation was performed with the argon laser at 488 nm while the emitted fluorescence of GFP and mKate was recorded at 484–536 nm and 567–654 nm, respectively. For each competition, three images from three independent seedlings were obtained. Representative images were used to visualize root colonization. Zen 3.1 Software (Carl Zeiss) and ImageJ was used for image visualization.

### Genome re-sequencing and genome analysis of single isolates and endpoint populations

Genomic DNA of selected evolved isolates from different time points and whole populations from the final transfer (T30) of the EE was extracted from 2 mL ON cultures using the EURx Bacterial and Yeast Genomic DNA Kit. Resequencing of single evolved isolates was performed as previously described (Dragoš et al., 2018b, 2018c; Gallegos-Monterrosa et al., 2021; Martin et al., 2020; Thérien et al., 2020). Briefly, paired-end libraries were prepared using the NEBNext® Ultra™ II DNA Library Prep Kit for Illumina. Paired-end fragment reads were generated on an Illumina NextSeq sequencer using TG NextSeq® 500/550 High Output Kit v2 (300 cycles). Primary data analysis (base-calling) was carried out with “bcl2fastq” software (v2.17.1.14, Illumina). All further analysis steps were done in CLC Genomics Workbench. Reads were quality-trimmed using an error probability of 0.05 (Q13) as the threshold. In addition, the first ten bases of each read were removed. Reads that displayed ≥80% similarity to the reference over ≥80% of their read lengths were used in the mapping. Non-specific reads were randomly placed to one of their possible genomic locations. Quality-based SNP and small In/Del variant calling was carried out requiring ≥8 × read coverage with ≥25% variant frequency. Only variants supported by good quality bases (Q ≥ 20) were considered and only when they were supported by evidence from both DNA strands in comparison to the *B. subtilis* NCIB 3610 genome and pBS plasmid (GenBank accession no. NZ_CP020102 and NZ_CP020103, respectively).

For whole-population sequencing of the evolved populations from the final transfer, acoustic fragmentation PCR-free libraries were prepared using MGIEasy PCR-free Library Prep Set (MGI Tech). Paired-end fragment reads (150bp x 2) were generated on a DNBSEQ-Tx sequencer (MGI Tech) following the manufacturer’s procedures. All population samples were sequenced with >200 x coverage for polymorphism calling. Raw data were filtered using SOAPnuke (v1.5.6) (Chen et al., 2018) with parameters “-l 15 -q 0.08 -n 0.03 -T 1 -Q 2 -5 0 -G -M 2” to remove low quality reads (more than 8% of bases with quality lower than 15, N content more than 3%). Mutations were called using *breseq* (v0.35.7) with the default parameters and a -p option for population samples (Deatherage and Barrick, 2014). The default parameters called mutations only if they appeared at least 2 times from each strand and reached a frequency of at least 5% in the population. Similar to single evolved isolates, the *B. subtilis* NCIB 3610 genome and pBS plasmid were used as references for mutation calling. For both single evolved isolates and whole populations, mutations were removed if they were also found in the ancestor to obtain the final mutation set. Identified mutations in the evolved isolates and whole populations are listed in Table S1 and S2, respectively.

The sequencing data for single evolved isolates that support the findings of this study has been deposited into the NCBI Sequence Read Archive (SRA) database under BioProject accession number: PRJNA705352, and sequencing data for endpoint populations into CNGB Sequence Archive (CNSA) (Guo et al., 2020) of China National GeneBank DataBase (CNGBdb) (Chen et al., 2020) with accession number CNP0002416.

## QUANTIFICATION AND STATISTICAL ANALYSIS

### Data and statistical analysis

Data and statistical analysis were carried out in Excel, OriginPro and R Studio. Outliers were identified using Dixon’s test for outliers. For all statistical tests, normality was evaluated with a Shapiro-Wilk test. Equal variance was tested using F-test (for two groups) or Levene’s test (for more than two groups). In addition, P-values were corrected for multiple testing using the Benjamini & Hochberg (BH) procedure within each series of experiments. For all statistical tests, a significance level of 0.05 was used. No statistical methods were used to pre-estimate sample size and the experiments were not randomized.

For individual root colonization, the log10-transformed productivity (CFU/mm root) of the replicates were divided by the mean of the log10-transformed productivity of the ancestor from the same experimental setup. For competition between WT and the Δ*hag* mutant, the observed frequency of the WT replicates after 48 h were divided by 0.5 (the starting frequency in the inoculation mix). For these experiments, the resulting normalized values were subjected to a One-sample *t*-test to test whether the mean was significantly different from 1. For pairwise competitions between the ancestor and evolved isolates on the root or in LB + xylan, the relative fitness (r) of the evolved isolate was calculated by comparing the frequency of the evolved isolate at the beginning and the end of the competition experiment as shown in equation 1 (Jousset et al., 2009; Li et al., 2021a; Ross-Gillespie et al., 2007), in which X_0_ is the initial and X_1_ is the final frequency of the evolved isolate. The relative fitness was log2-transformed, and these values were subjected to a One-sample *t*-test to test whether the mean was significantly different from 0.

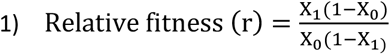

For root co-colonization of the ancestor and Ev7.3 with the community, within each inoculation ratio, the difference between *B. subtilis* ancestor and Ev7.3, and between the communities co-inoculated with the ancestor and with Ev7.3, was tested with a Two-sample *t*-test. If the groups had unequal variance, Welch’s Two-sample *t*-test was applied.

For swimming and swarming motility, an ANOVA was performed on the log10-transformed data at each time point followed by a Dunnett’s Multiple Comparison test using the ancestor as the control group.

For growth curve analysis of the ancestor and evolved isolates in monoculture, the carrying capacity (K) of individual replicates was calculated from the OD_600_ data using the Growthcurver-package in R (Sprouffske and Wagner, 2016). Significant difference in carrying capacity between ancestor and evolved isolates was tested by an ANOVA followed by a Dunnett’s Multiple Comparison test. Similarly, for the growth profiles of *B. subtilis* ancestor and Ev7.3 in co-culture with the community, the carrying capacity (K) was calculated from the GFP_485/20nm_ data. Significant difference between ancestor and Ev7.3 under the same inoculation ratio or alone was tested by a Two-sample *t*-test or Wilcoxon Unpaired Two-sample test (when data failed to meet parametric assumptions).

