## Supplementary material for "Experimental evolution of *Bacillus subtilis* on *Arabidopsis thaliana* roots reveals fast adaptation and improved root colonization in the presence of soil microbes": Fig S1 to S7 and Table S1

### Supplementary Figures

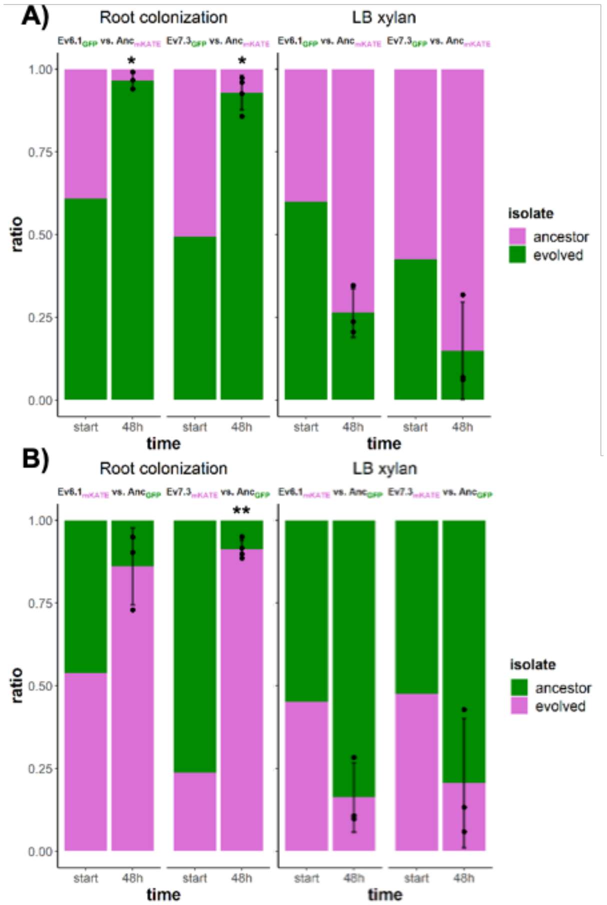


**Fig. S1:** Competition between selected evolved isolates and the ancestor during root colonization and in LB xylan under shaking conditions. (**A**) Competition between ancestor (magenta) and evolved isolates (green) during root colonization (left) and in LB xylan (0.5 %) under shaking conditions (right) for 48 h. (**B**) Same as in A, but with opposite fluorescent labels. The bar plots show the starting ratio of the evolved isolate and ancestor in the mix, and the observed ratios after 48 hours. Bars represent the mean (N=3-4) and the error bars represent standard deviation. The points show the replicates for the evolved (from below) and ancestor (from above). For statistical analysis, the relative fitness (r) of the evolved isolates was calculated by comparing the frequency of the evolved isolate at the beginning and at the end of the competition experiment. The log2-transformed relative fitness values were subjected to a One-sample *t*-test to test whether the mean was significantly different from 0. * and ** indicate $P<0.05$ and $P<0.01$, respectively.

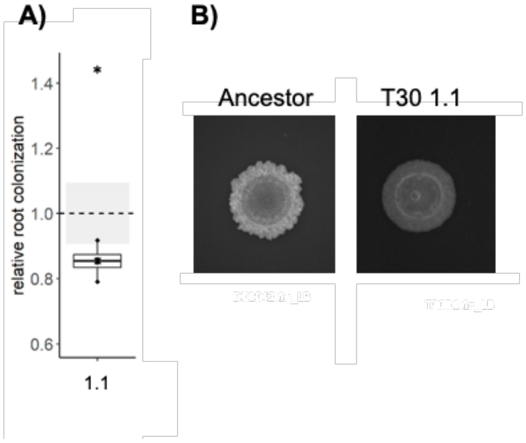


**Fig. S2:** Relative root colonization and colony morphology of an evolved isolate from population 1 at the final transfer. (**A**) Isolate Ev1.1 from transfer 30 was tested for individual root colonization. Relative root colonization was calculated by dividing the log10-transformed productivity (CFU/mm root) of each replicate by the mean of the log10-transformed productivity of the ancestor. The cross represents the mean relative root colonization (N=4). The dashed, horizontal line represents the mean of the ancestor (N=4), while the grey-shaded rectangles represent the standard deviation of the ancestor. The normalized values were subjected to a One-sample *t*-test to test whether the mean was significantly different from 1. * indicates $P<0.05$. (**B**) ON cultures of the ancestor and Ev1.1 were spotted on LB agar (1.5 %) and imaged after incubation for 48 h at 30 °C using the stereomicroscope. Ancestor represents *B. subtilis* DK1042. Each colony is representative of at least three replicates. Scale bar denotes 5 mm.

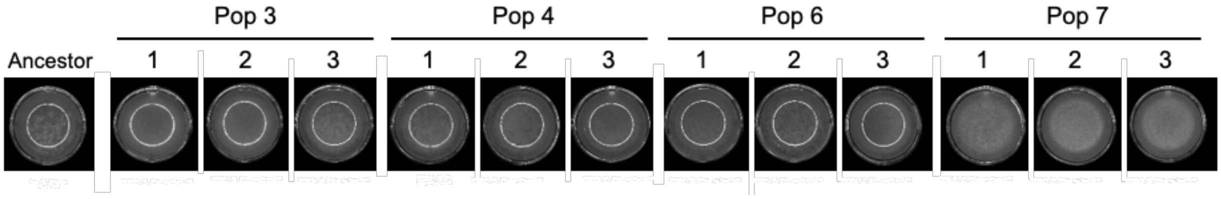


**Fig. S3:** Pellicle biofilm formation in LB of the ancestor and evolved isolates from the final transfer. *B. subtilis* ancestor and evolved isolates from transfer 30 were inoculated into LB medium at a starting $OD_{600}$ of 0.05 in 24-well plates and incubated for 48 h at 30 °C. Each image is representative of 3-4 replicates.

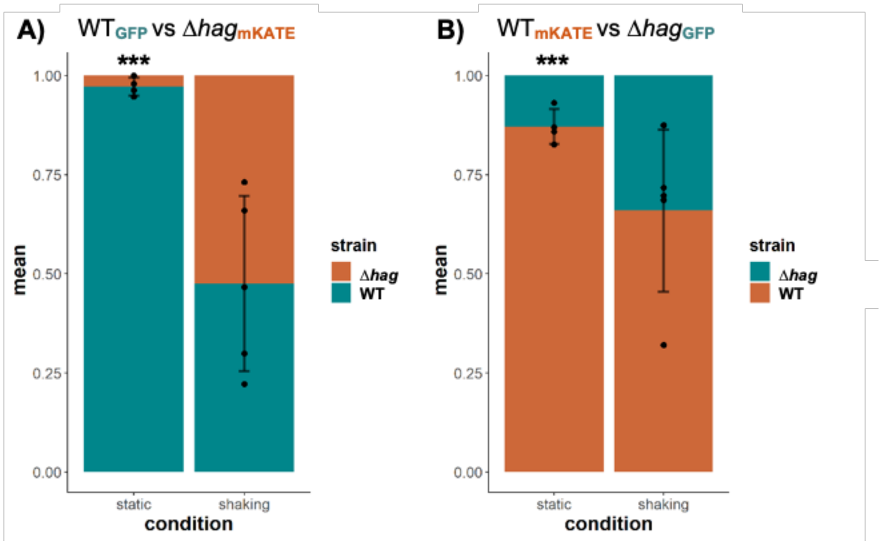


**Fig. S4:** Motility is not important for root colonization under shaking conditions. Competition between *B. subtilis* DK1042 (WT) and a Δ*hag* mutant during root colonization in MSNg medium. *A. thaliana* seedlings were inoculated with a 1:1 mix of WT and mutant with opposite fluorescent labels at a starting $OD_{600}$=0.02. Plates were incubated for 48 h in the plant chamber under static conditions or shaking conditions (200 rpm). After three successive rounds of root colonization, productivity on the third root was quantified as CFU per cm root. (**A**) Competition between WT (turquoise) and the Δ*hag* mutant (orange). (**B**) Same as in A, but with opposite fluorescent labels. Bars represent the mean ratio (N=4-5) and the error bars represent standard deviation. The points show the replicates for the WT (from below) and the Δ*hag* mutant (from above). The observed frequency of the WT replicates after 48 h were divided by 0.5 (the starting frequency in the inoculation mix), and the resulting normalized values were subjected to a One-sample *t*-test to test whether the mean was significantly different from 1. *** indicates P<0.001.

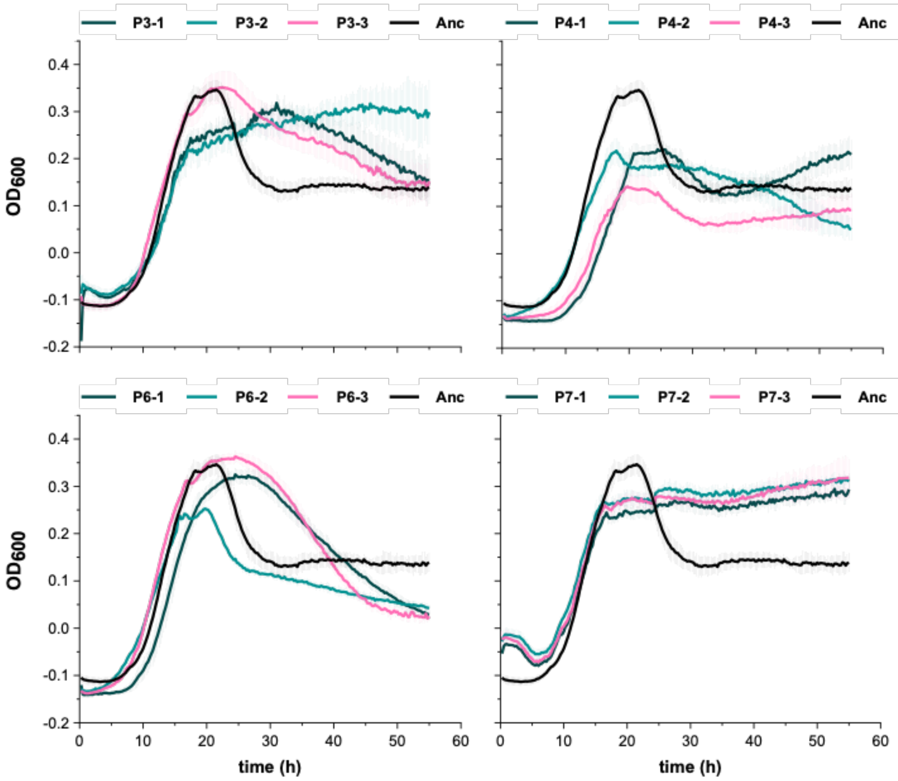


| Carrying capacity (K) | | |
|---|---|---|
| **Isolate** | **Estimate** | **P-value** |
| **3.1** | 0.06171 | <0.001 *** |
| **3.2** | 0.1046 | <0.001 *** |
| **3.3** | 0.05412 | <0.001 *** |
| **4.1** | -0.01126 | 0.868 |
| **4.2** | -0.03859 | <0.001 *** |
| **4.3** | -0.0955 | <0.001 *** |
| **6.1** | 0.01045 | 0.912 |
| **6.2** | -0.06691 | <0.001 *** |
| **6.3** | 0.01917 | 0.254 |
| **7.1** | 0.07988 | <0.001 *** |
| **7.2** | 0.10695 | <0.001 *** |
| **7.3** | 0.09901 | <0.001 *** |

**Fig. S5:** Growth of *B. subtilis* ancestor and evolved isolates in MSNc + xylan. Growth of the ancestor and evolved isolates in MSNc + xylan (0.5 %) (starting $OD_{600}$ = 0.1) was measured every 15 min at 24 °C under shaking conditions (orbital). Data represents mean and error bars represent standard deviation (N=6-12, 2 independent ON cultures with 3-6 technical replicates each). The carrying capacity (K) was calculated using "Growthcurver" in R. Significant difference between ancestor and evolved isolates was tested by an ANOVA followed by a Dunnett's Multiple Comparison test. *** indicates P<0.001.

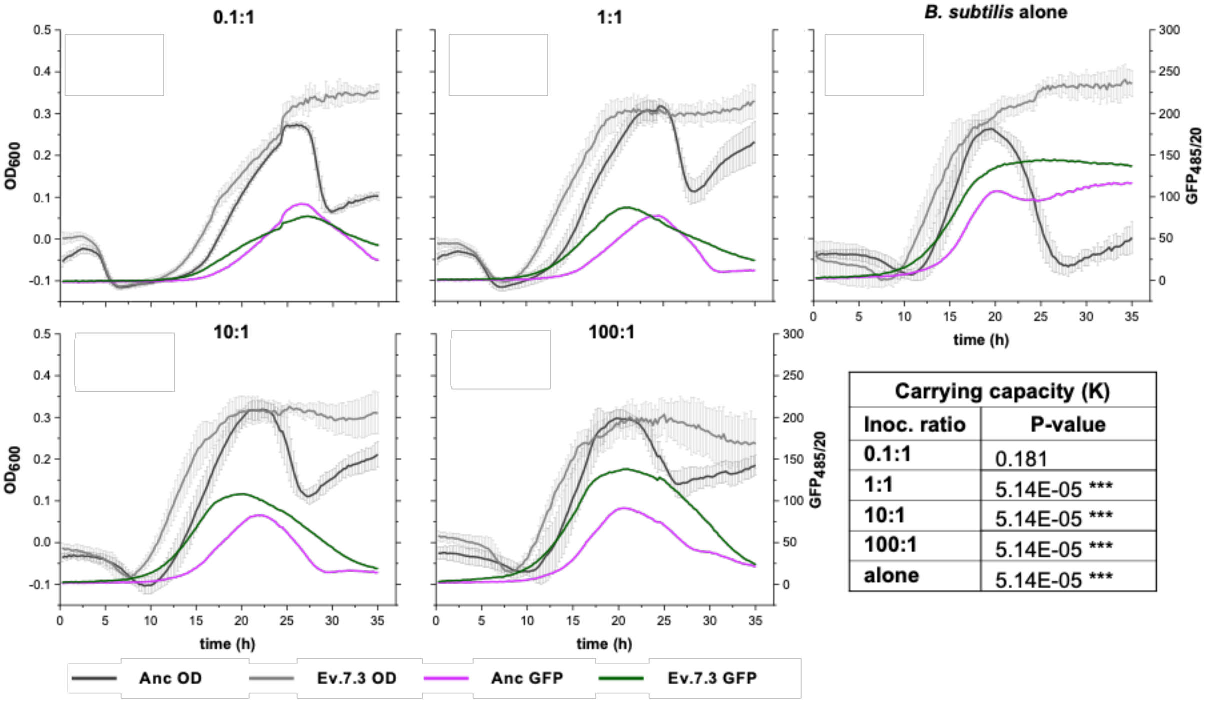


| Carrying capacity (K) | |
|---|---|
| **Inoc. ratio** | **P-value** |
| **0.1:1** | 0.181 |
| **1:1** | 5.14E-05 *** |
| **10:1** | 5.14E-05 *** |
| **100:1** | 5.14E-05 *** |
| **alone** | 5.14E-05 *** |

**Fig. S6:** Ev7.3 shows enhanced carrying capacity in MSNc + xylan when co-cultured with a semi-synthetic, soil-derived community compared to the ancestor. Growth of constitutively GFP-expressing *B. subtilis* ancestor or Ev7-3 in co-culture with the community in MSNc + xylan (0.5 %) was measured under four different inoculation ratios: 0.1:1, 1:1, 10:1 and 100:1 of *B. subtilis* and community, respectively. $OD_{600}$ and $GFP_{485/20nm}$ were measured every 15 min for 35 h at 24 °C while shaking. Data represents mean and error bars represent standard deviation (N=6-9, 2-3 independent ON cultures with 3 technical replicates each). The carrying capacity (K) was calculated from the $GFP_{485/20nm}$ data. Significant difference in carrying capacity between the ancestor and Ev7.3 under the same inoculation ratio or alone was tested by a Two-sample *t*-test or Wilcoxon Unpaired Two-sample test (when data failed to meet parametric assumptions). *** indicates P<0.001.

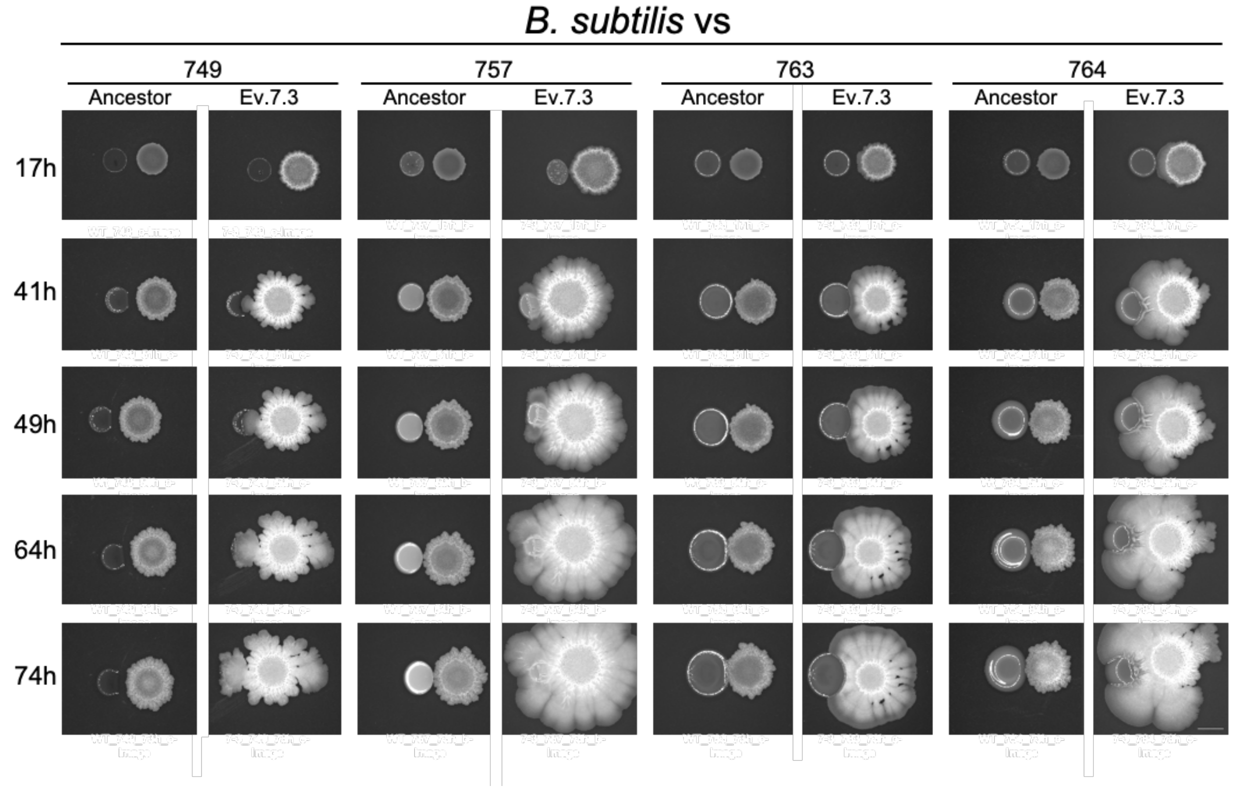


**Fig. S7:** Pairwise interactions of *B. subtilis* ancestor or evolved isolate 7.3 with bacterial species. Ancestor or Ev7.3 was spotted on LB agar (1.5 %) at 0.7 cm distance from bacterial species belonging to *Pedobacter sp.* D749, *Rhodococcus globerulus* D757, *Stenotrophomas indicatrix* D763 or *Chryseobacterium sp.* D764. Plates were incubated at 30 °C and images were captured at the given time points using the stereomicroscope. Ancestor represents *B. subtilis* DK1042. Each colony is a representative of three replicates. Scale bar denotes 5 mm.

**Table S1:** The table shows detected mutations in the re-sequenced genomes of evolved isolates. The functions of the gene products were retrieved from the SubtiWiki Database (Zhu and Stülke, 2018). T = Transfer.

| T12 | | | T18 | | | T30 | | | | | | | Position | Gene | Mutation | Function |
|---|---|---|---|---|---|---|---|---|---|---|---|---|---|---|---|---|
| **7.1** | **7.2** | **7.3** | **7.1** | **7.2** | **7.3** | **1.1** | **6.1** | **6.2** | **6.3** | **7.1** | **7.2** | **7.3** | | | | |
| | | | | | | | | X | | | | | 72039 | *spoIIE* | Glu489* | protein serine phosphatase, |
| | | | | | | | | | | | X | | 84980 | *pabC* | Leu24Val | aminodeoxychorismate lyase, |
| | | | | | | | | | | X | | X | 235527 | Intergenic (*glpT*/*ybeF*) | T>C | *glpT*: glycerol-3-phosphate permease. *ybeF*: unknown |
| | X | | | | | | | | | | | | 529455 | *tRNA* | C>T | |
| | | X | | | | | | | | | | | 578493 | *ydeS* | Ala39Val | unknown regulator (similar to transcriptional regulator (TetR family)) |
| | | X | | | | | | | | | | | 961914 | *ssuB* | 483C>T | aliphatic sulfonate ABC transporter |
| | | | | | | X | | | | | | | 1002530 | Intergenic (*glpP*/*glpF*) | G>T | *glpP*: transcriptional antiterminator, regulation of glycerol and glycerol-3-phosphate utilization. *glpF*: glycerol facilitator, glycerol uptake |
| | | | | | | | X | X | X | | | | 1002601 | *glpF* | Cys22_Val25del | glycerol facilitator, glycerol uptake |
| | | X | | | | X | | | | | | | 1073693 | *scoC* | Ala21Asp | transition state regulator |
| | | | | | | | X | | | | | | 1221482 | *oppA* | Thr534Asn | oligopeptide ABC transporter |
| | | | | | | | X | | X | | | | 1471759 | *kinA* | Gly568Val | two-component sensor kinase, initiation of sporulation |
| | | | | | | | | | | | | X | 1472932 | *patA* | 138G>A | aminotransferase, biosynthesis of lysine and peptidoglycan |
| | | X | | | | | | | | | | | 1528830 | *pdhA* | 474G>T | pyruvate dehydrogenase, links glycolysis and TCA cycle |
| X | | | | | | | | | | | | | 1567847 | *ylbD* | p.Glu56* | outer spore coat protein, |
| | | | | | | X | | | | | | | 1573820 | *ddcP* | *342Leu | DNA damage checkpoint recovery protease |
| | | | | | | X | | | | | | | 1686014 | *trmFO* | Arg58Ser | tRNA:m(5)U-54 methyltransferase, tRNA modification |
| | | | | | | | X | | X | | | | 1702691 | *fliM* | p.Arg326Ile | flagellar motor switch protein, movement and chemotaxis |
| | | | | | | | X | | X | | | | 1947405 | *galU* | 672C>T | UTP--glucose-1-phosphate uridylyltransferase |
| | | | | | | | | | | X | | | 2026710 | *yoaE* | p.Val427Leu | formate dehydrogenase |

| | | | | | | | | | | | | | | | | |
|---|---|---|---|---|---|---|---|---|---|---|---|---|---|---|---|---|
| | | | | | | X | | | | | | | 2406541 | *ypbD* | His116Asp | unknown |
| | | | | | | X | | | | | | | 2414116 | *rsiX* | Lys200fs | anti-SigX, control of SigX activity |
| | | | X | | | | | | | | | | 2422584 | *spmB* | p.Ala81Ser | spore maturation protein (spore core dehydratation) |
| | | | | | | | | | X | | | | 2434346 | *ypuB* | p.Glu9* | hypothetical (unknown) |
| X | | | X | X | X | | | | | X | X | X | 2552672 | Intergenic (*sinI*/*sinR*) | C>G | sinI: antagonist of SinR<br>sinR: transcriptional regulator, control of biofilm formation |
| | | | | | | | | | | X | | | 2759185 | *sacC* | Arg304fs | levanase, degradation of levan to fructose |
| | | | | | | X | | | | | | | 2893562 | *leuA* | 150C>A | 2-isopropylmalate synthase, biosynthesis of leucine |
| | | | X | | | | | | | | | | 3005087 | *tcyN* | Gly19Val | cystine ABC transporter (ATP-binding protein), cystine uptake |
| | | | | | | | | | | | | X | 3070742 | *ythQ* | Glu226Asp | *ythQ* = function unknown, similar to ABC transporter |
| | | | | | | | | | | X | | X | 3141769 | *ythB* | 948G>A | cytochrome bd2, menaquinol oxidase, respiration |
| | | | | | | | | | | X | | X | 3409591 | *sigO* | Met139del | RNA polymerase sigma factor |
| | | | X | X | | | | | | | | | 3498891 | *yvfR* | Arg127Ile | ABC transporter (ATP-binding protein) |
| | | X | | | | | | | | | | | 3537728 | *levB* | His70Tyr | endolevanase, levan degradation |
| | | | | X | | | | | | | | | 3547528 | *pgmB (pgcM?)* | Asp14Tyr | beta-phosphoglucomutase, starch and maltodextrin utilization |
| | | X | | | | | | | | | | | 3570846 | *glmR* | Pro223His | regulator of carbon partitioning between central metabolism and peptidoglycan biosynthesis |
| | | | | | | X | | | | | | | 3666013 | *gtaB* | Cys116Phe | UTP-glucose-1-phosphate uridylyltransferase, biosynthesis of teichoic acid |
| | | | | | | X | | | | | | | 3666017 | *gtaB* | A>G | |
| | | | | | | X | | | | | | | 3666019 | *gtaB* | .Arg118Pro | |
| | | | | | | X | | | | | | | 3666048 | *gtaB* | Val128Leu | |
| | | | | | | X | | | | | | | 3666050 | *gtaB* | A>T | |
| | | | | | | X | | | | | | | 3666059 | *gtaB* | T>A | |
| | | | | | | X | | | | | | | 3666080 | *gtaB* | Glu138Asp | |
| | | | | | | X | | | | | | | 3672992 | *yvzE* | Leu2Arg | |

| | | | | | | | | | | | | | | | | |
|---|---|---|---|---|---|---|---|---|---|---|---|---|---|---|---|---|
| | | | | | | X | | | | | | | 3673014 | *yvzE* | 27T>C | putative UTP-glucose-1-phosphate uridylyltransferase |
| | | | | | | | | X | | | | | 3679471 | *tagE* | His330Tyr | UDP-glucose:polyglycerol phosphate glucosyltransferase, biosynthesis of teichoic acid |
| | | | X | X | | | | | | | | | 3679534 | *tagE* | Glu309* | |
| | | | | | X | | | | | X | X | X | 3679795 | *tagE* | Glu222* | |
| | | | | | | | X | | X | | | | 3679856 | *tagE* | Trp202fs | |
| | | | | | | X | | | | | | | 4025538 | *wapA* | Asn1683fs | cell wall-associated protein precursor, intercellular competition |
| | | | | | | X | | | | | | | 4151147 | *walH* | Arg252Thr | negative effector of WalK, controls cell wall metabolism |
| | | X | | | | | | | | | | | 4152067 | *walK* | Asp554Tyr | two-component sensor kinase, control of cell wall metabolism |
| | | | | | | | X | X | X | | | | 4152909 | *walK* | Thr273Lys | |
